# Toward a comprehensive modification landscape of yeast mitochondrial tRNAs using Nanopore direct RNA sequencing and Dihydrouridine sequencing

**DOI:** 10.1101/2025.05.09.653160

**Authors:** Julia L. Reinsch, David M. Garcia

## Abstract

*Saccharomyces cerevisiae* is an invaluable model in the study of mitochondrial tRNA biology. Yet the positions of modified bases in all yeast mitochondrially-encoded tRNAs (mt-tRNAs) are still not fully mapped. We performed Nanopore direct RNA sequencing (DRS) on tRNAs from the crude mitochondrial fraction of yeast to map base modifications across all 24 mt-tRNA isoacceptors. Additionally, we developed a method to detect dihydrouridine sites in tRNAs, tD-seq, where chemical reduction of dihydrouridine causes disruptions to reverse transcription. We mapped dihydrouridine, pseudouridine, and N2-dimethylguanosine sites in mt-tRNAs using DRS, tD-seq, and knockouts of five conserved tRNA-modifying enzymes. Our results establish Dus1 and Dus2 as the enzymes responsible for D_14_, D_16_, D_17_, D_17a_, and D_20_ formation in *S. cerevisiae* mt-tRNAs, and revealed interactions between Dus1, Dus2, and Trm1-catalyzed modifications. We provide a comprehensive analysis of *S. cerevisiae* mt-tRNA base modifications, and identify novel modification “circuits” in yeast mt-tRNAs, in which the loss of a single enzyme’s activity can change modification levels at sites catalyzed by other enzymes. These findings expand our understanding of mt-tRNA base modifications and their interdependence, and advance opportunities for the yeast model for investigating defects in human mt-tRNA function.

**GRAPHICAL ABSTRACT:** 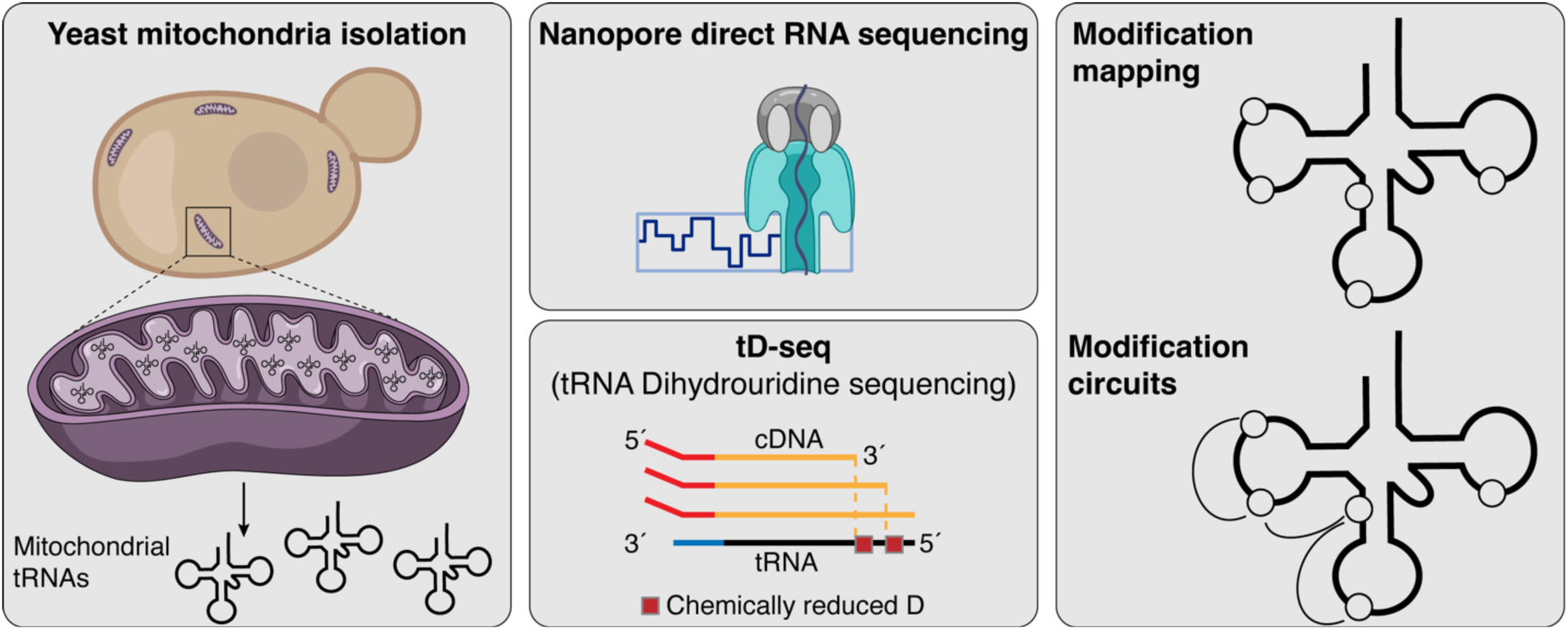

## INTRODUCTION

Eukaryotic organisms contain a mitochondrial genome in the form of circular dsDNA molecules found in the mitochondrial matrix. The yeast mitochondrial genome is a highly reduced derivative of its prokaryotic ancestor (1), yet it retains genes involved in oxidative phosphorylation that are essential for respiration. These genes are transcribed, then translated within mitochondria by translation machinery that includes mitochondrially encoded rRNA and tRNA. Mitochondrial tRNAs (“mt-tRNAs”) are chemically modified, though typically to a lesser extent than cytosolic tRNAs (2). These chemical modifications serve important roles in tRNA structure, stability, and promotion of translation fidelity (reviewed in (3)).

Pseudouridine and dihydrouridine are the two most abundant tRNA base chemical modifications in nature. Pseudouridine (Ψ) is an isomer of uridine that augments its hydrogen bonding and base stacking properties (4). Pseudouridine synthases (Pus) catalyze the formation of Ψ at positions 27, 28, 31, 32, 38, 39, 55, and 72 in mt-tRNAs (5). Dihydrouridine (D) is a chemically reduced form of uridine that alters the planarity of the base, conferring flexibility to the D-loop (6) and increasing the overall stability of tRNA tertiary structure (7). While D is found at positions 14, 16, 17, and 20 in mt-tRNAs (8), the enzymes responsible for D formation in the mitochondria have not been previously identified.

Disruption to the chemical modifications of mt-tRNAs can lead to defects in cell growth and are responsible for several human diseases, referred to as “modopathies” (9, 10). Comprehensive annotation of human mt-tRNA modifications was performed through analysis of all 22 individually purified human mt-tRNAs by LC-MS/MS (11), demonstrating an important approach to mapping base modifications, albeit with a high technical benchmark. This approach is less amenable to investigating the impact of multiple cell or enzyme perturbations on modifications simultaneously across the majority of isoacceptors, highlighting the need for another approach.

The budding yeast *S. cerevisiae* has been a powerful model system for investigating the enzymes that chemically modify mt-tRNA for several reasons: 1) Yeast genetics make it straightforward to perturb modification enzymes; 2) Yeast can survive by fermentation, making mitochondrial gene expression non-essential under appropriate growth conditions; 3) Many of the tRNA modifications and the enzymes that catalyze them are conserved in human mitochondria; 4) Yeast cultures can be easily scaled in order to obtain sufficient material for analysis. Discoveries enabled by work in yeast have helped develop a mechanistic understanding of the basis of several human modopathies (12–15). Nevertheless, characterization of yeast mt-tRNA modifications is incomplete, with the principal modification database, Modomics, providing modification profiles for only 16 of 24 yeast mt-tRNA isoacceptors (8). A complete annotation of all mt-tRNA modifications in yeast will further our understanding of their role in mitochondrial function and disease.

We used Oxford Nanopore Technologies (ONT) direct RNA sequencing (DRS) of full length tRNAs to analyze tRNA chemical modifications. This measurement can provide position-level resolution at increased throughput compared to mass spectrometric analysis. As an RNA molecule transitions through a protein-based nanopore, its chemically modified bases can lead to distinct ionic current signals compared to unmodified bases. These ionic current changes can cause basecalling programs to artifactually assign a base to the modified position that does not match the “reference”, unmodified base. This outcome is referred to as a base “miscall”, and certain modifications can produce distinct miscall patterns. For example, pseudouridine typically causes a U-to-C miscall coincident with the modified position. Coupled with biochemical and/or genetic controls, miscalls are informative about the positions and identities of modified bases. These approaches have been used to study base modifications in conditions that affect the cellular growth state or RNA modifying enzyme activities (16–21). Additionally, DRS can provide an *in vivo* systems-level understanding of modification interdependencies across dozens of tRNAs (20)

Using RNA from the crude mitochondrial fraction of budding yeast cell lysate, we analyzed yeast mt-tRNAs by DRS. We built our DRS-based sequence analysis on a revised complete structure-guided alignment of *S. cerevisiae* mt-tRNA sequences. Using the Modomics database as a reference, we found that DRS can detect nearly all known yeast mt-tRNA modifications, some supported using genetic mutants. We orthogonally validated all dihydrouridine sites in mt-tRNAs, using a method that we developed, tD-seq, which led to identification of new examples of modification interdependencies. Our study extends the utility of DRS for analysis of tRNA sequences and modifications, and presents a new method to determine the positions of dihydrouridine in tRNA. This work also advances this model for investigating the mechanistic basis of human modopathies caused by defects in tRNA chemical modifications.

## RESULTS

### A revised structural alignment of all 24 *S. cerevisiae* mitochondrial tRNAs

To analyze the sequences and modifications of yeast mt-tRNAs, we first generated an updated reference list of mt-tRNA sequences and a corresponding structural alignment. Prior to our study, the Modomics database contained 16 mt-tRNA sequences from *S. cerevisiae*, yet its mitochondrial DNA encodes at least 24 mt-tRNA genes (22). The eight isoacceptors not currently in the Modomics database are: mt-tRNA^Ala(UGC)^, mt-tRNA^Asn(GUU)^, mt-tRNA^Asp(GUC)^, mt-tRNA^Cys(GCA)^, mt-tRNA^Glu(UUC)^, mt-tRNA^Gln(UUG)^, mt-tRNA^Val(UAC)^. To ensure the accuracy of our reference sequences, we performed pairwise comparisons of mt-tRNA sequences annotated from RNA-seq experiments (21), Modomics (8), and NCBI (23, 24). While many mt-tRNA sequences were identical among these sources, we found that 19 of the 24 mt-tRNAs in the NCBI database contained additional nucleotides at the 5′ and/or 3′ends. We removed these extra nucleotides from our reference since they are not predicted to form additional base pairs at the ends of the acceptor stem and were outside the bounds determined by RNA-seq. The resulting mt-tRNA sequence reference is summarized in Supplementary Table S1. It includes the RNA bases of the 5′ splint adapter used in our sequencing experiments followed by the mature mt-tRNA sequence, and then the RNA bases of the 3′ splint adapter. We focused our analysis on the tRNA sequences encoded in the mitochondrial genome, and did not attempt to quantify the abundance of tRNAs imported into the mitochondria from the cytosol (25). This was because our enrichment protocol did not stringently purify mitochondrial material from the cytosol, and thus it was not possible to distinguish nuclear-encoded tRNAs that were imported into the mitochondria, from the same molecules present in the cytosolic fraction.

Next, we manually curated a structure-guided sequence alignment of all 24 mt-tRNAs based on conserved tRNA structural features including loop length, stem length, and tertiary contacts (Figure 1A). We adopted a conventional tRNA numbering scheme (26), with accommodations for predicted single nucleotide bulges and additional nucleotides in the D-loop and anticodon-loop. Apart from these minor differences, our predicted secondary structures were overall very similar to those in our previous analysis of cytosolic tRNAs (20). As seen in the cytosolic tRNAs, the Serine and Leucine mt-isoacceptors are both of the “type II tRNAs”, containing longer variable loops (27). In mitochondria, the Tyrosine tRNA also has a longer variable loop, unlike the cytosolic Tyrosine tRNA. No mt-tRNAs had predicted secondary structures dramatically different from a canonical tRNA, such as a missing D-arm or unusually long variable regions, as observed in some animals and archaea (reviewed in (28)). We used this structural alignment and numbering scheme in the analysis and presentation of our sequencing data (Figure 1A). We summarize the modifications found in yeast mt-tRNAs and their catalyzing enzymes, if known (Figure 1B).

**Figure 1.**
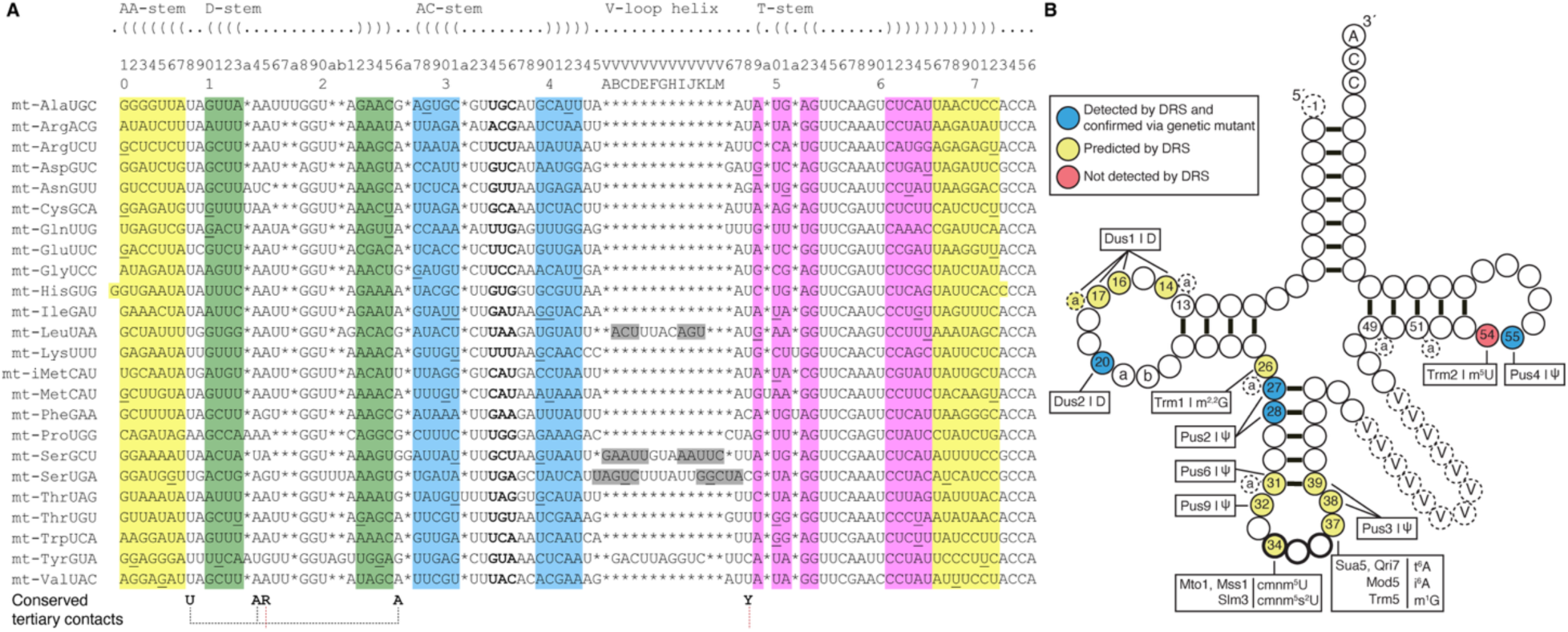
Budding yeast mitochondrial tRNA (mt-tRNA) sequences and the ensemble of their modifications profiled in this study. (**A**) Structure-guided sequence alignment of 24 mt-tRNA isoacceptors from *S. cerevisiae*. Each base position of the alignment matches a box in the heatmaps presented in Figures 3–5. Asterisks indicate gaps. Left and right parentheses and colored regions show predicted helices/stems (labeled at top). Numbers (top line) indicate each base position in a repeating series of ∼ten, or (below) every ∼tenth position. The variable loop is up to 13 base positions in length, and alignment of these bases, as well as the bases at neighboring positions (44–48), in the Leucine, Serine and Tyrosine isoacceptors, were anchored via predicted helices/stems in the loop. Histidine tRNA contains a “-1” G added post-transcriptionally (8). G•U wobbles are included in predicted stem regions and underlined. Conserved tertiary contacts are indicated underneath the alignment (R=purine, Y=pyrimidine). (**B**) Secondary structure-based illustration of a generic yeast mt-tRNA, highlighting overall structural segments, ten types of known chemical modifications, and the names of single enzymes or heteromeric complexes annotated to catalyze them. Positions in blue are those that contain modifications verified in this study by their corresponding genetic mutants. Positions in yellow are those that also trigger base miscalls and are predicted to be chemically modified bases in at least one isoacceptor in this study. Position 54 (red) is annotated to be modified to m^5^U in many mt-tRNAs, but this modification does not yield a base miscall coincident with this position, as previously documented (20).

### Direct sequencing of all 24 *S. cerevisiae* mitochondrial tRNAs

In our previous sequencing of total yeast tRNA isolated from wild-type cells grown in glucose and harvested during exponential phase (20), the abundance of reads aligning to our mt-tRNA reference was 0.3% of total, while the remaining 99.7% mapped to a reference set of 42 cytosolic tRNA isoacceptor sequences. Two factors resulted in low representation of mt-tRNA: 1) mitochondrial tRNAs are significantly less abundant than cytosolic tRNAs, and 2) yeast cells grown in media containing glucose as a principle carbon source contain fewer mitochondria than cells grown in carbon sources that require oxidative respiration, such as glycerol or ethanol (29). To address these limitations, we grew yeast cells in large cultures (1L) of 2% ethanol media and performed an enrichment for yeast mitochondria prior to RNA isolation (Figure 2A).

**Figure 2.**
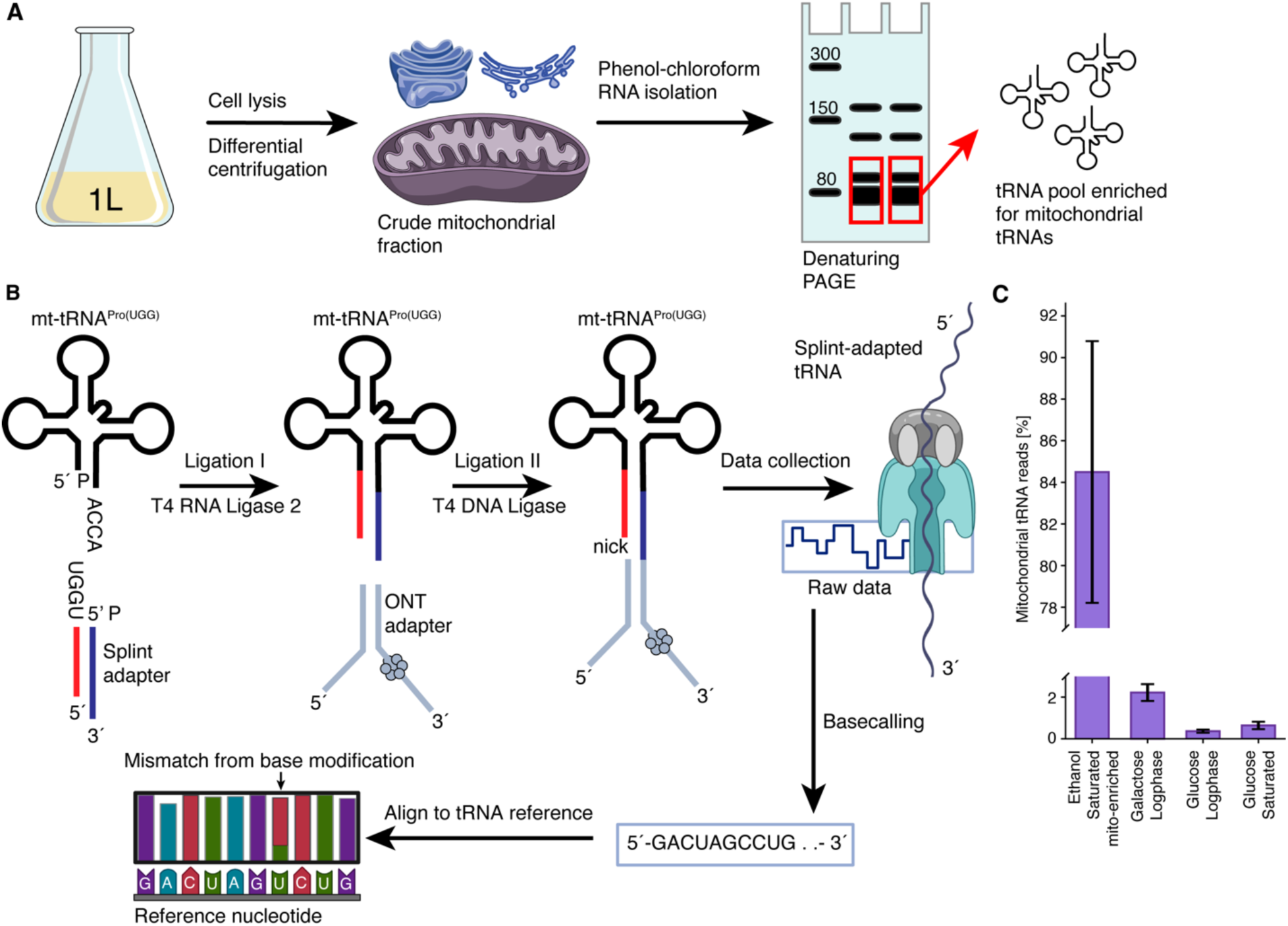
Overview of sample preparation and tRNA library preparation and sequencing. (**A**) Crude mitochondrial fraction from large yeast cultures is subjected to traditional RNA isolation, followed by gel purification of tRNA fraction. (**B**) tRNAs are ligated to a double-stranded splint adapter with T4 RNA Ligase 2 by the tRNA’s 3′ NCCA overhang. A second ligation is performed using T4 DNA Ligase with the tRNA and ONT sequencing adapters. The splint-adapted tRNA pool is sequenced and based called as described in Materials and Methods. Base miscalls are analyzed to infer presence of chemical base modifications, using an IVT mt-tRNA library as a background model. (**C**) The full length mt-tRNA fraction from wild-type cells, as measured by DRS, was enriched ∼300-fold in comparison with a results obtained with total cellular tRNA isolated from cells grown on a fermentable carbon source (20).

We isolated RNA molecules between ∼70 and 100 nt long from the mitochondrially-enriched RNA fraction using PAGE. Oligonucleotide splint adapters were ligated to this RNA pool, followed by ligation to the Oxford Nanopore Technologies (ONT) RNA sequencing adapter (Figure 2B). The library of adapted tRNA molecules were sequenced with ONT MinION R9.4.1 flow cells and SQK-RNA002 kits. As a control for these cellular-derived tRNA molecules, we analyzed *in vitro* transcribed (IVT) mt-tRNA sequences that contained no modifications and thus served as a reference for DRS signal obtained from otherwise identical tRNA sequences (See Materials and Methods for details about IVT sample, library preparation, and sequencing). The raw ionic current patterns were converted to nucleotide base calls using ONT Guppy software (version 3.0.3) (Figure 2B). Each tRNA DRS experiment yielded between 34, 843 and 112, 800 total reads per flow cell (Supplementary Table S3).

We aligned reads to our curated reference containing all 24 mitochondrial and 42 cytosolic yeast tRNA isoacceptors (Supplementary Table S1) using BWA-MEM (30). Each tRNA DRS experiment yielded between 15, 337 and 63, 709 MAPQ1-aligned tRNA reads per flow cell (Supplementary Table S3). The average percentage of mapped reads across experiments was ∼51%, which is consistent with prior tRNA DRS experiments using RNA002 kits (Lucas et al. 2023; Shaw et al. 2024). Between 11, 904 and 62, 516 MAPQ1-aligned tRNA reads per flow cell aligned to mitochondrial tRNA reference sequences (Supplementary Table S3). This corresponds to between 78 and 98% of all tRNA reads, a substantial increase in mt-tRNA representation (∼300-fold above reference (20)) compared to prior tRNA DRS studies performed without mitochondrial enrichment (20, 21) and DRS of mitochondrial RNA without tRNA-specific capture (31) (Figure 2C).

For these analyses, we used the now-discontinued R9.4.1 flow cells and RNA002 sequencing kits because 1) data collection began prior to the release of ONT “RNA” flow cells and RNA004 kits, and 2) to permit direct comparison to our recent DRS analysis of 42 yeast cytosolic tRNAs using the same chemistry and basecalling approach. The miscall-based modification detection method, using the RNA002 chemistry employed here, was validated by orthogonal LC-MS/MS experiments in our prior study (20). Miscall-based DRS analysis of tRNA chemical modifications has provided valuable biological insights in numerous studies (16–20, 32) and will continue to do so until direct ionic current analysis of DRS data is robust and widely adopted.

### Direct RNA Sequencing reveals modification landscape of yeast mt-tRNAs

We generated heatmaps displaying reference match probabilities at each position in all 24 yeast mt-tRNAs to visualize base miscalls. The ‘reference match probability’ (RMP) at a given position in a tRNA represents the probability that the nucleotide assigned by the base caller matches the canonical unmodified reference nucleotide. These probabilities were calculated using an error-based model that accounts for standard DRS rates of mismatches, insertions, and deletions, as well as sequence context-based effects unique to the unmodified sequences that were derived from our IVT mt-tRNA data and other parameters (see Materials and Methods). The mt-tRNA IVT heatmap shows that nearly all positions have a high RMP (dark blue) as expected from unmodified RNA (Figure 3A). The IVT sequences represent an important control dataset that accounts for DRS error that can arise from certain sequence contexts of unmodified RNA bases. For example, while position 42 in mt-tRNA^Asp(GUC)^ here yielded a light-colored square — frequently associated with a modification-triggered miscall — we verified that the DNA oligonucleotide used as the template for *in vitro* transcription of this tRNA matched the reference nucleotide (G), using Sanger sequencing (data not shown). Further inspection of the k-mer associated with this position, 5′-**G**GAGG-3′, in other IVT molecules revealed a similar G-to-A miscall (20), implicating this as a sequence context-dependent miscall. Miscalls caused only by sequence context were rare across the other 1800+ sequenced nucleotide positions (Figure 3A).

**Figure 3.**
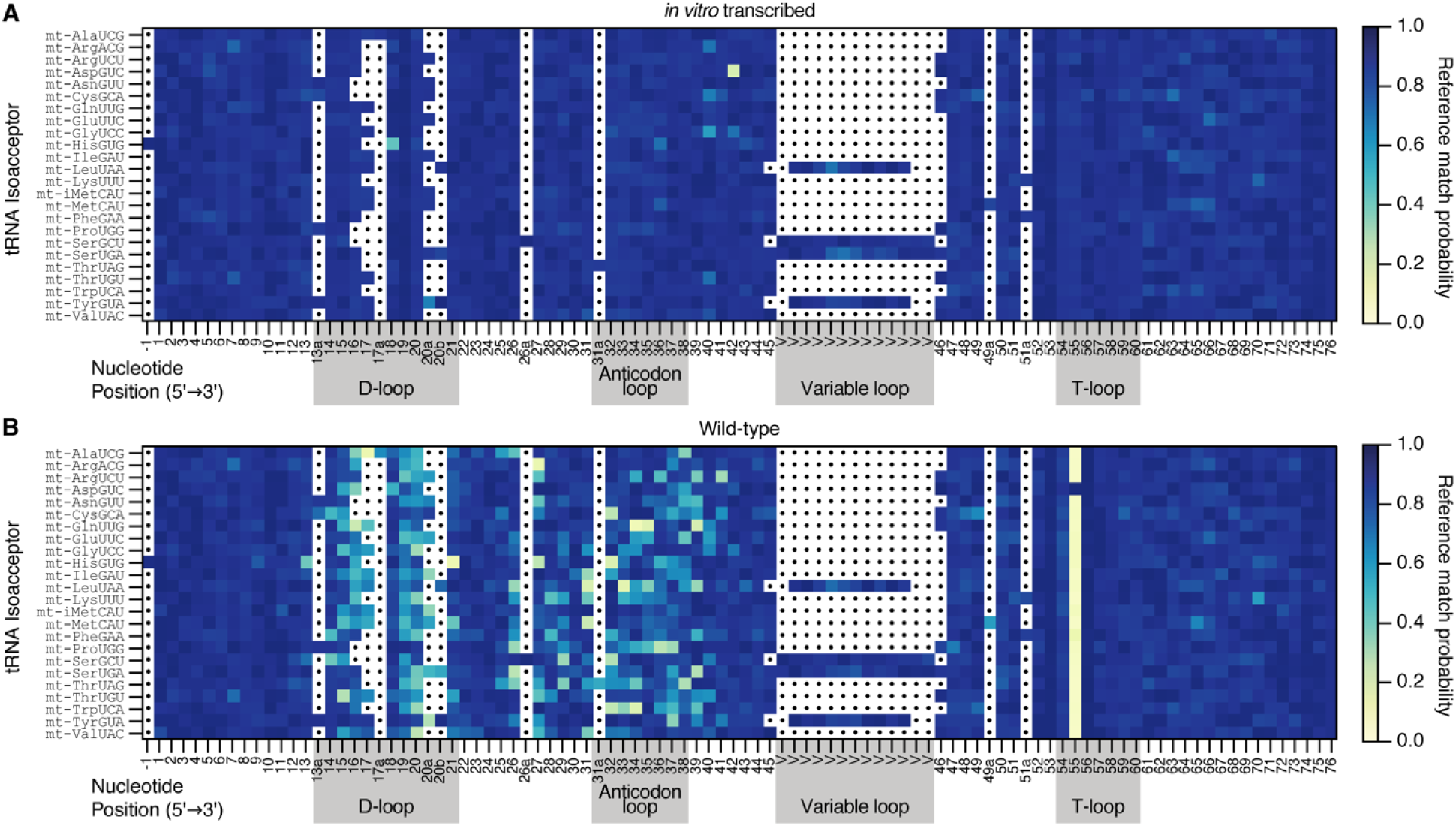
Heatmaps representing alignments of 24 *S. cerevisiae* mt-tRNA isoacceptors exhibit miscalls coincident with chemically modified positions. (**A**) Heatmap representing *in vitro* transcribed (IVT) tRNA sequences of 24 *S. cerevisiae* mitochondrial isoacceptors. A higher reference match probability (dark blue) corresponds to positions where the base-called nucleotide more frequently concurs with the reference nucleotide given the alignment, and a lower reference match probability (light yellow) corresponds to positions where the base-called nucleotide more frequently disagrees with the reference nucleotide. Sequences were aligned by the D-loop, anticodon loop, variable loop, and T-loop (grey boxes). Positions are numbered according to a conventional tRNA base numbering scheme, based on the sequence alignment in Figure 1A. (**B**) Wild-type isoacceptors isolated from yeast cells. Otherwise as described above.

In contrast to the IVT data, the heatmap of wild-type mt-tRNAs isolated from yeast, shows many positions with low RMPs (yellow-toned) spanning the D-loop to the T-loop even after correction from the error model, indicating a high frequency of base miscalls and possible modification sites (Figure 3B). There are notably fewer base miscalls in these mitochondrial tRNAs relative to cytosolic tRNAs analyzed with the same method (20), in agreement with a recent literature review (3).

Within the 16 previously annotated mt-tRNAs present in the Modomics database, 10 unique chemical modifications were documented in 16 different base positions — some modifications appear at more than one position, such as pseudouridine (Supplementary Table S4) (8). We calculated the *mismatch* probability (equal to 1.0 minus the reference match probability value) for each position from a random subset of 105 aligned reads per mt-tRNA isoacceptor. In the program used to calculate posterior probabilities, marginCaller, the default posterior probability threshold to call an alternative nucleotide was empirically determined and set at 0.3 (33). Positions with a mismatch probability <0.3 in IVT and ≥0.3 in wild-type were classified as true miscalls. Of the 16 documented modification sites, 15 had a mismatch probability of ≥0.3 in at least one mt-tRNA annotated to have a modification at that position (Supplementary Table S5). The only annotated modification that was not detected on any mt-tRNA isoacceptor by DRS was m^5^U_54_, a result consistent with our prior sequencing of cytosolic yeast tRNAs, in which genetic deletion of the enzyme that catalyzes this modification (Trm2) did not alter the DRS-based signal at U_54_ (20). Therefore, as the vast majority of annotated modifications in mt-tRNAs (15 out of 16) produced a detectable miscall-based signal with our protocol, these data highlight the utility of DRS for profiling tRNA modifications. Given the strong correlation between mt-tRNA modifications as documented on Modomics with miscall-based signals in our DRS data, we extrapolated to the other eight mt-tRNAs whose ensemble of modifications were not previously documented. We predict these contain approximately 5–8 modifications each, a density resembling that of the other 16 isoacceptors. An exception was the sparsely modified mt-tRNA^Asp(GUC)^ that only produced miscalls at 2 positions (Supplementary Table S5). Most of these eight mt-tRNAs likely also contain Trm2-catalyzed m^5^U_54_, as 15 of 16 mt-tRNA isoacceptors listed in the Modomics database contain this modification (8).

Among known mt-tRNA modifications, pseudouridine was most robustly detected by miscalls in our DRS data. Pseudouridine can be detected as a U-to-C miscall at the modified position and does not typically impact miscalls at neighboring, unmodified positions (34). In wild-type cells, Ψ sites at positions 27, 28, 31, and 55 were recorded as miscalls in 100% of previously annotated positions, while Ψ sites at positions 32 and 39 were recorded as miscalls in 5 of 6 isoacceptors previously annotated to contain them (Supplementary Table S4). The only previously annotated Ψ position that our data did not recapitulate was position 72 of mt-tRNA^iMet(CAU).^ We considered the possibility that this was due to differences in growth conditions as our cells were grown in ethanol as a carbon source while the original study used galactose (35). Our analysis of mt-tRNA^iMet(CAU)^ grown in galactose, however, did not produce a miscall at position 72 (Supplementary Figure S1). As the original study noted the possibility of “incomplete modification of U_72_ to Ψ”, it remains possible that a low fraction of modified molecules preclude detection by DRS. Finally, we identified 21 sites likely to be pseudouridylated in the eight unannotated mt-tRNAs, based on comparison to Ψ positions known in the 16 annotated mt-tRNAs. We subsequently validated 11 of these 21 sites through genetic knockouts of pseudouridine synthases.

### Pus4 pseudouridylates U_55_ in 23 of 24 mt-tRNAs in *S. cerevisiae*

The Pus4 enzyme catalyzes formation of U_55_ to Ψ_55_ in 41 of 42 tRNA isoacceptors in the yeast cytosol (20). Pus4 also modifies mt-tRNAs (36) and all 16 of the yeast mt-tRNAs with known sequences and modification profiles contain Ψ_55_ (8). We sequenced tRNAs from the crude mitochondrial fraction of yeast lacking the *PUS4* gene to verify that the low reference match probability at U_55_ shown in the heatmap of wild-type mt-tRNAs (Figure 3B) was due to Pus4-catalyzed Ψ. In mt-tRNAs isolated from this *pus4*Δ strain, there was a high reference match probability at position 55 (Figure 4A), indicating a lack of modification. The subtractive heatmap highlights differences in reference match probability between mt-tRNA isolated from wild-type and *pus4*Δ yeast strains, where blue-colored squares represent a decrease in modification in the mutant (Figure 4B). Out of the 24 yeast mt-tRNAs, we found that 23 were modified at position 55 by Pus4 (Supplementary Table S6). The exception was mt-tRNA^Asp(GUC)^ which has a guanosine at position 55 and therefore cannot be pseudouridylated.

**Figure 4.**
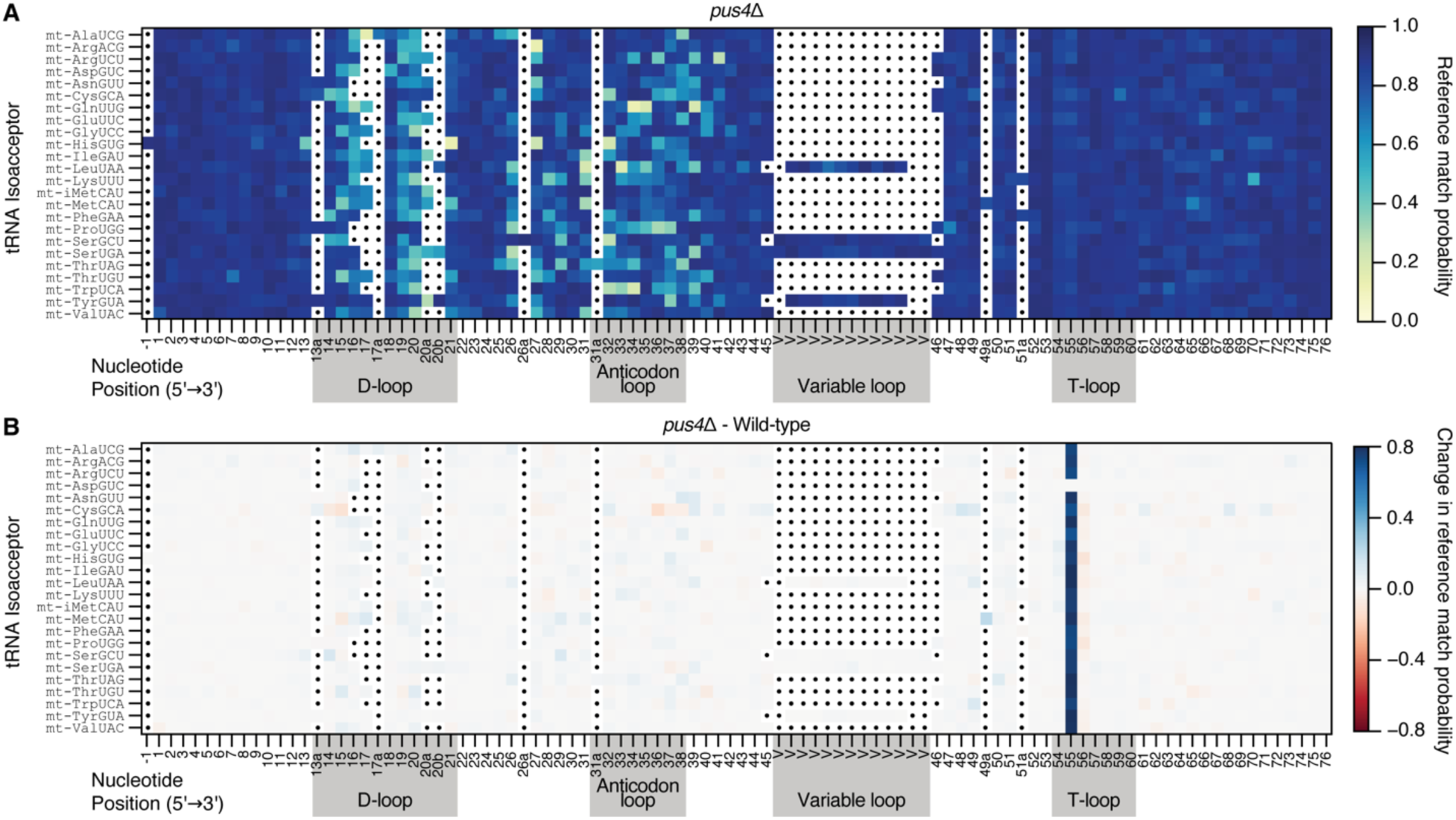
Heatmaps demonstrate Pus4-based pseudouridylation of 23 of 24 yeast mt-tRNA isoacceptors. (**A**) Heatmap representing mt-tRNAs from *pus4*Δ cells. Otherwise as described in Figure 3. (**B**) The change in reference match probabilities between *pus4*Δ and wild-type aligned isoacceptors. A positive change in reference match probability (blue squares) indicates a decrease in base miscalls for *pus4*Δ, and a negative change in reference match probability (red squares) indicates an increase in base miscalls for *pus4*Δ. White-toned squares indicate that reference match probabilities do not differ between the two strains. The scale was determined by the maximum change in reference match probability.

Some tRNA modifications are known to be involved in “circuits” where one modification promotes or represses the formation of other modifications (37). A well-studied modification circuit involves Pus4-catalyzed Ψ_55_ promoting addition of m^1^A_58_ in the T-loop of many yeast cytosolic tRNAs (20, 38, 39). Notably, reduction of m^1^A_58_ levels in cytosolic tRNAs upon the loss of Ψ_55_ has been observed by DRS (20). The enzyme complex catalyzing the formation of m^1^A_58_ in cytosolic tRNAs — Trm6 and Trm61 — is not known to localize to budding yeast mitochondria (40, 41), and no budding yeast mt-tRNAs are annotated to contain this modification (8). Our data support the view that the m^1^A_58_ does not occur in any mt-tRNAs, as we did not detect miscalls associated with m^1^A_58_ in 24 mt-tRNAs. Furthermore, we did not find evidence of a modification circuit initiated by Pus4, as there were no additional changes beyond the 55^th^ position in *pus4*Δ cells (Figure 4B). Our inability to detect m^5^U_54_ by DRS precludes evaluation of whether Pus4 influences this modification in mt-tRNAs, as has been shown to occur in cytosolic tRNAs (20, 38).

### Identification of Pus2-dependent Ψ sites in *S. cerevisiae* mt-tRNAs

In budding yeast, Pus2 is a mitochondrially-localized enzyme that modifies U_27_ and U_28_ of mt-tRNAs, while its paralog Pus1 modifies cytosolic tRNAs at the same positions (42, 43). Seven of the 16 yeast mt-tRNAs currently in the Modomics database are annotated to contain Ψ at position 27 or 28 (8), but only sites on mt-tRNA^Arg(UCU)^ and mt-tRNA^Lys(UUU)^ have been determined to be Pus2-dependent (43). We mapped Pus2-dependent modification sites on all 24 mt-tRNAs by sequencing tRNA isolated from the crude mitochondrial fraction of yeast with a deletion of the *PUS2* gene, (Figure 5A, Supplementary Figure S2A), and we assigned Pus2-dependent sites by comparing the mismatch probabilities between *pus2*Δ and wild-type (Supplementary Table S5). We classified Ψ sites as Pus2-dependent if they met the following criteria: 1) the mismatch probability was equal to or greater than the 0.3 threshold in the wild-type data, 2) the mismatch probability was less than 0.3 in the *pus2*Δ data, and 3) the difference in these mismatch probabilities met or exceeded an empirically determined minimum difference threshold (see Materials and Methods). This analysis corroborated all seven previously annotated Ψ_27_/Ψ_28_ sites across different mt-tRNAs and showed that Pus2 modifies six of these sites (Supplementary Table S6). The exception was U_27_ of mt-tRNA^Thr(UAG)^, where the mismatch probability was above 0.3 at position 27 in both wild-type and *pus2*Δ. This miscall is likely caused by Ψ as previously annotated (44), but we cannot conclude that Pus2 is the specific pseudouridine synthase that modifies mt-tRNA^Thr(UAG)^ at this site. Additionally, we inferred the presence of a Pus2-dependent Ψ_27_ for three mt-tRNAs that have not been previously annotated: mt-tRNA^Asn(GUU)^, mt-tRNA^Thr(UGU)^, and mt-tRNA^Val(UAC)^ (Supplementary Table S6). For mt-tRNA^Thr(UGU)^, we also found evidence for a Pus2-dependent miscall at U_28_, which would make it the only mt-tRNA to have Ψ at both positions 27 and 28, a feature that is known to exist in Pus1-catalyzed sites in cytosolic tRNAs (e.g. in tRNA^Lys(UUU)^, tRNA^Val(CAC)^ and tRNA^Trp(CCA)^)(8, 20).

**Figure 5.**
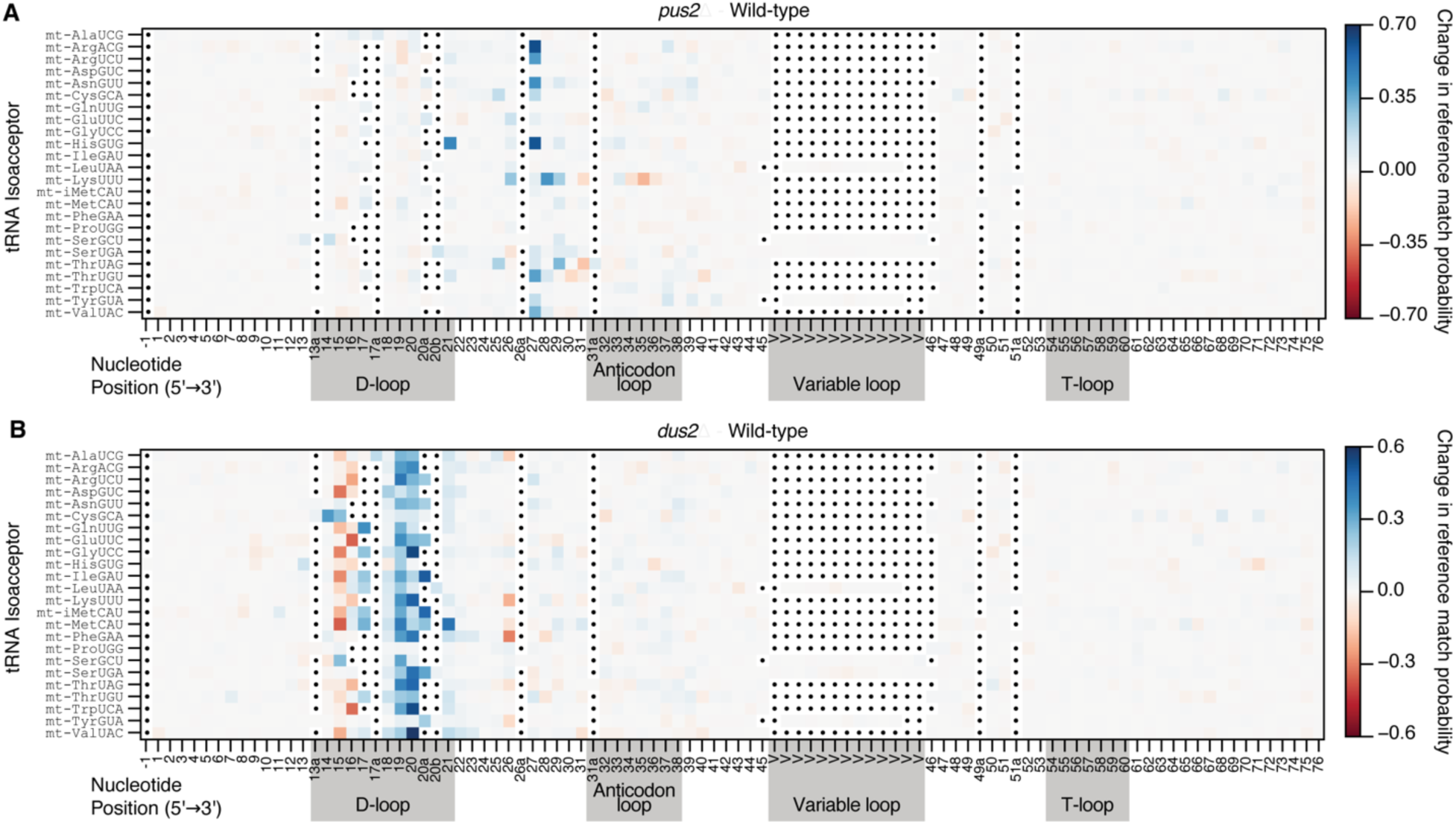
Heatmaps reveal both positions with modifications catalyzed by Pus2 and Dus2 and other impacted sites. The heatmaps show the change in reference match probabilities between (**A**) mt-tRNAs from *pus2*Δ and wild-type cells, and (**B**) mt-tRNAs from *dus2*Δ and wild-type cells. Otherwise as described in Figure 4B. The scales were determined by the maximum change in reference match probability.

### Pus2-catalyzed Ψ_27_ and Ψ_28_ impact miscalls at other modification sites

Our sequencing of mt-tRNA from *pus2*Δ yeast revealed that loss of Ψ_27_/Ψ_28_ induced changes in base miscalls at other sites. These changes suggest Pus2 is involved in a modification “circuit”, an instance in which one modification promotes or represses addition of a second, different modification (37, 45). In some cases, loss of Ψ_27_/Ψ_28_ led to an associated *increase* in miscalls, or putative modification, at a different site. We observed this pattern in mt-tRNA^Lys(UUU)^, mt-tRNA^Thr(UAG)^, and mt-tRNA^Thr(UGU)^, where the reference match probability was decreased at position 31 or 30, when cells were missing Pus2. Position 31 is annotated as Ψ in mt-tRNA^Thr(UAG)^ and mt-tRNA^Lys(UUU)^ (8), therefore the modification levels of Ψ_31_ may be increased in *pus2*Δ for these isoacceptors. However, position 30 of mt-tRNA^Thr(UGU)^ is a guanosine, which is a position not known to be modified in any yeast tRNA. Therefore, the source of this signal is unknown. For mt-tRNA^Lys(UUU)^, the reference match probability at positions 34 and 35 were lower in *pus2*Δ compared to wild-type, suggesting that loss of Ψ_28_ leads to an increase in a modification elsewhere. We initially hypothesized that this change was caused by an increase in levels of the annotated modification cmnm^5^s^2^U at position 34 of the mt-tRNA^Lys(UUU)^ anticodon. A heterodimeric complex composed of Mto1 and Mss1 forms cmnm^5^U while Slm3 independently forms s^2^U at position 34 of mt-tRNAs (12). We used APM-PAGE followed by Northern Blotting to measure 2-thiolation levels of mt-tRNA^Lys(UUU)^ in wild-type and *pus2*Δ strains (Supplementary Figure S3A, B).

The results indicated that the loss of Ψ_28_ does not significantly alter 2-thiolation levels of mt-tRNA^Lys(UUU)^. It is possible that the increase in miscalls was due to increased levels of the non-thiolated portion of this modified uridine (cmnm^5^U), which is present in the anticodon of tRNA^Lys(UUU)^ at a relatively low stoichiometry in wild-type cells (12). However, cmnm^5^U is not detectable by APM-PAGE. Alternatively, the change in miscall could have been caused by the addition of a different modification not previously annotated at position 34/35 in this isoacceptor. It is unclear what impact a change in modification at this position would have on the ability of mt-tRNA^Lys(UUU)^ to decode mRNA codons, although cmnm^5^U_34_, s^2^U_34_, and cmnm^5^s^2^U_34_ all promote mt-tRNA^Lys(UUU)^ charging and mitochondrial protein synthesis (12, 46).

Additionally, we found a putative modification circuit in which one modification promotes the formation of a subsequent modification. In mt-tRNA^His(GUG)^, there was a large increase in reference match probability at A_21_ in *pus2*Δ relative to wild-type (Figure 5A), which suggests that the addition of Ψ_27_ by Pus2 may promote the formation of an unknown modification at or near A_21_ in wild-type cells. The miscall at A_21_ was not dependent on the presence of D_20_ (see below) and therefore cannot be attributed to a change in dihydrouridine levels. Moreover, this site did not cause stops or misincorporations during reverse transcription in our tD-seq data from wild-type samples (see below), suggesting that it is not a modified adenosine that interferes with processivity of the SuperScript III reverse transcriptase. Since there are no modifications known to occur at A_21_ in any tRNAs in any organism, further experiments are needed to identify the source of this signal.

### Identification of Dus2 sites in *S. cerevisiae* mt-tRNAs by DRS

All 16 mt-tRNA isoacceptors currently in the Modomics database are annotated to contain dihydrouridine (D) at position 20, yet the enzyme that catalyzes this modification has not been confirmed experimentally. It has long been presumed that Dus2, which is responsible for dihydrouridylating position 20 in yeast cytosolic tRNAs (47), also modifies mitochondrial tRNAs at the same position. This presumption is supported by the detection of Dus2 in the yeast mitochondrial proteome (40, 41). Our DRS analyses of mt-tRNAs from *dus2*Δ yeast provide experimental evidence that Dus2 is, in fact, necessary for D_20_ formation in mt-tRNAs. In *dus2*Δ cells, we observed a decrease in modification at position 20 and/or neighboring positions 19 and 20a/20b/21 for many mt-tRNAs (Figure 5B, blue squares; Supplementary Figure S2B). Therefore, a three-position window, centered on position 20, was considered for our analysis of mismatch probabilities, such that a threshold change at any one or more of these positions lead us to assign a D_20_ (see Materials and Methods for details). While the miscall signature of dihydrouridine has not been previously examined in the context of a genetic mutant, this three-position window has been observed for other genetically validated modifications, such as m^1^A_58_ (20). Of the 16 mt-tRNAs in the Modomics database annotated to contain D_20_, DRS corroborated 13 sites as Dus2 targets (Supplementary Table S6), indicating that DRS can detect dihydrouridine in this context. The three mt-tRNAs annotated to contain D_20_ sites that were not detected by DRS were: mt-tRNA^Pro(UGG)^, mt-tRNA^His(GUG)^, and mt-tRNA^Tyr(GUA)^. In these three tRNAs, the modification was detectable in these isoacceptors by an orthogonal, D-specific method as Dus2-dependent (see below). Of the eight mt-tRNAs not currently in the Modomics database, DRS detected Dus2-dependent changes in six and did not detect D_20_ in mt-tRNA^Asp(GUC)^ and mt-tRNA^Cys(GCA)^, though both contain a uridine at position 20.

### Identification of all D sites by tRNA Dihydrouridine sequencing (tD-seq)

Above, we analyzed dihydrouridine using DRS supported by data from a genetic knockout of a dihydrouridine synthase. In addition to changes that we observed coincident with the annotated modification site of Dus2 (U_20_), we observed unexpected and significant changes at positions 15/16 in many mt-tRNAs (Figure 5B). As dihydrouridine occurs at other positions in the D-loop that could be impacted by loss of Dus2, we sought to clarify our DRS-based predictions of D sites in mt-tRNAs using an orthogonal method. ‘D-seq’ (48, 49) and ‘Rho-seq’ (50) were developed to map D-sites across the nuclear-encoded transcriptome in *S. cerevisiae* and *S. pombe,* respectively. These methods use sodium borohydride (NaBH_4_) to chemically reduce dihydrouridine into tetrahydrouridine (51), which may either cause cDNA terminations (RT stops) one nucleotide 3′of the D site (48, 50), or base misincorporations coincident with a D site (52), during reverse transcription. While some cytosolic tRNA reads were present in the libraries of these previous studies, the presence of modifications that produce NaBH_4_-*independent* RT-stops 3′ of D-loop sites — such as m^2, 2^G_26_, m^1^G_37_, and m^1^A_58_ — precluded analysis of most D sites in these tRNAs. Additionally, neither study was designed to specifically capture tRNAs nor enrich for mitochondrial RNAs.

To address this gap, we developed a new method called ‘tD-seq’ (tRNA <u>D</u>ihydrouridine <u>seq</u>uencing) to map dihydrouridine sites in mt-tRNAs. The chemistry of this protocol was based on the original ‘D-seq’ protocol (49), but the library preparation method was modified to enrich tRNAs (Figure 6A). Due to the lower frequency of the aforementioned common RT-stop modifications in mt-tRNAs relative to cytosolic tRNAs, this method yields sufficient coverage of the D-loop of nearly all mt-tRNA isoacceptors without demethylation or other treatment. In samples from wild-type yeast not treated with NaBH_4_ (Supplementary Figure 4A), RT stops occurred at or adjacent to established m^2, 2^G_26_ and m^1^G_37_ sites, which impair cDNA synthesis (53). Upon NaBH_4_ treatment of wild-type RNA (Figure 6B and Supplementary Figure S4B), RT stops appeared at several positions in the D-loop of most isoacceptors, as anticipated for this treatment, and unexpectedly within the anticodon loop of several isoacceptors where D is not known to exist. In contrast, the combined knockout of *DUS1* and *DUS2—*which encode the only D-synthases found in yeast mitochondria (40, 41)*—*led to the loss of NaBH_4_-induced stops in the D-loop (Figure 6C and Supplementary Figure S4C). This result demonstrates that tD-seq can robustly detect dihydrouridine sites in tRNA.

**Figure 6.**
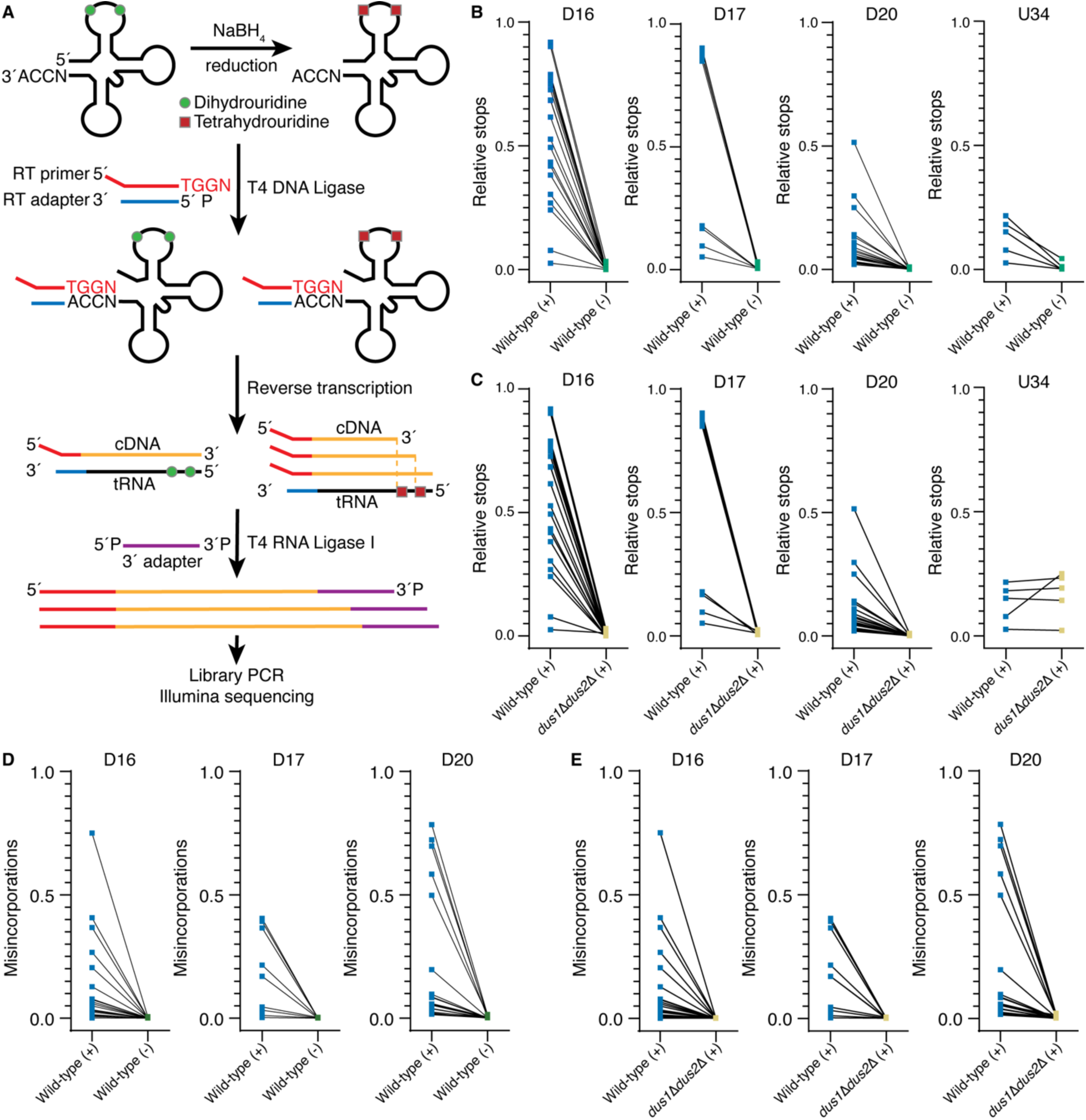
tD-seq detects dihydrouridine sites in mt-tRNAs. (**A**) Overview of tD-seq protocol, where RNA is treated with NaBH_4_ which converts dihydrouridine into tetrahydrouridine. A 3′RT primer is ligated to tRNAs with T4 DNA ligase prior to reverse transcription with SuperScript III RT. The 3′adapter is ligated to cDNA using T4 RNA Ligase I and then PCR is performed prior to Illumina sequencing. (**B**) RT stops induced by NaBH_4_ treatment, relative to the coverage of the position 1 nt 3′ of the specified site. Each square point represents the average relative stops for a single mt-tRNA isoacceptor at the specified site. Points from the same isoacceptor are connected by black lines. On the x-axis, NaBH_4_ treatment is specified with (+) or (-) next to the genotype. The position 17 and 34 plots have fewer isoacceptors because not all mt-tRNAs have a base and/or a uridine at these positions in the structural alignment. (**C**) NaBH_4_-induced stops in the D-loop of mt-tRNAs are absent in *dus1*Δ*dus2*Δ, while the stops at U34 are observed independently of Dus activity. (**D**) Misincorporation levels compared between matched wild-type samples with and without NaBH_4_ treatment. (**E**) Misincorporation levels compared between three biological replicates of wild-type and *dus1*Δ*dus2*Δ.

We next considered an explanation for the unanticipated NaBH_4_-induced RT stops within the anticodon loop. While D has never been found at these positions in any tRNA, the stop positions coincide with known cmnm^5^U_34_/cmnm^5^s^2^U_34_ sites. Besides dihydrouridine, NaBH_4_ is known to chemically reduce several modified bases including ac^4^C, m^7^G, and m^1^A (54, 55). Our tD-seq data provides evidence that cmnm^5^U and/or cmnm^5^s^2^U modifications may also be affected by NaBH_4_ treatment; therefore, this method could be used to detect these modifications. This can be tested in the future by performing tD-seq on deletion strains of Mss1/Mto1 and Slm3, which collectively form cmnm^5^s^2^U_34_.

While tD-seq produced robust stops for some D sites (D_14_, D_16_, D_17_), D_20_ sites yielded less robust stops. Base misincorporations coincident with D sites generally followed a similar positional-pattern to RT stops, when comparing wild-type (untreated), wild-type (treated), and *dus1*Δ*dus2*Δ (treated) samples (Figure 6D, E and Supplementary Figure S5A–C). Therefore, we considered the change in both RT stops *and* misincorporations (between matched treated and untreated wild-type samples) to facilitate our tD-seq D site annotations, along with assignment of their catalyzing enzymes (Dus1 or Dus2). Cutoff thresholds were empirically determined by comparing values from ‘D’ and ‘non-D Uridine’ sites from annotations in the Modomics database (see Materials and Methods for details on D-site calling).

We used tD-seq to map D-sites in all 24 mt-tRNAs and found that each tRNA contains between one and four dihydrouridines within their D-loops. Comparison of D-site annotations between the Modomics database, DRS, and tD-seq showed a high level of agreement (Supplementary Table S8). Several isoacceptors in the Modomics database (mt-tRNA^Gly(UCC)^, mt-tRNA^iMet(CAU)^, mt-tRNA^Leu(UAA)^, and mt-tRNA^Tyr(GUA)^) are annotated to have a D at position 16 and a ‘U’ at position 17, however our tD-seq results clearly indicated that both positions 16 and 17 are dihydrouridines (Supplementary Figure S4B). Earlier methods used to analyze mt-tRNA sequences (56) may have lacked the resolution needed to detect neighboring D sites. We examined D-sites that were detected by tD-seq, but not DRS, to determine if there was a common feature such as sequence context or stoichiometry between these sites that could explain the lack of DRS signal. There were no obvious differences in sequence context between D-sites detected and not detected by DRS (Supplementary Figure S6). Additionally, several sites not detected by DRS produced robust RT stops or misincorporations in tD-seq, suggesting that these sites are not low stoichiometry.

Dus1 modifies cytosolic tRNAs at positions 16 and 17 (47), and is found in the yeast mitochondrial proteome (40). Our tD-seq data demonstrated that Dus1 catalyzes modifications at four different positions in mt-tRNAs: D_14_, D_16_, D_17_, and D_17a_ (Figure 7A, B and Supplementary Figure S7A, C), establishing its mitochondrial activity and expanding the list of its target sites. The remaining Dus enzymes in yeast, Dus3 and Dus4, were not detectable in the mitochondrial proteome (40, 41), and our data provided no evidence of their canonical activities at positions 47 or 20a/b, respectively, in mt-tRNAs.

**Figure 7.**
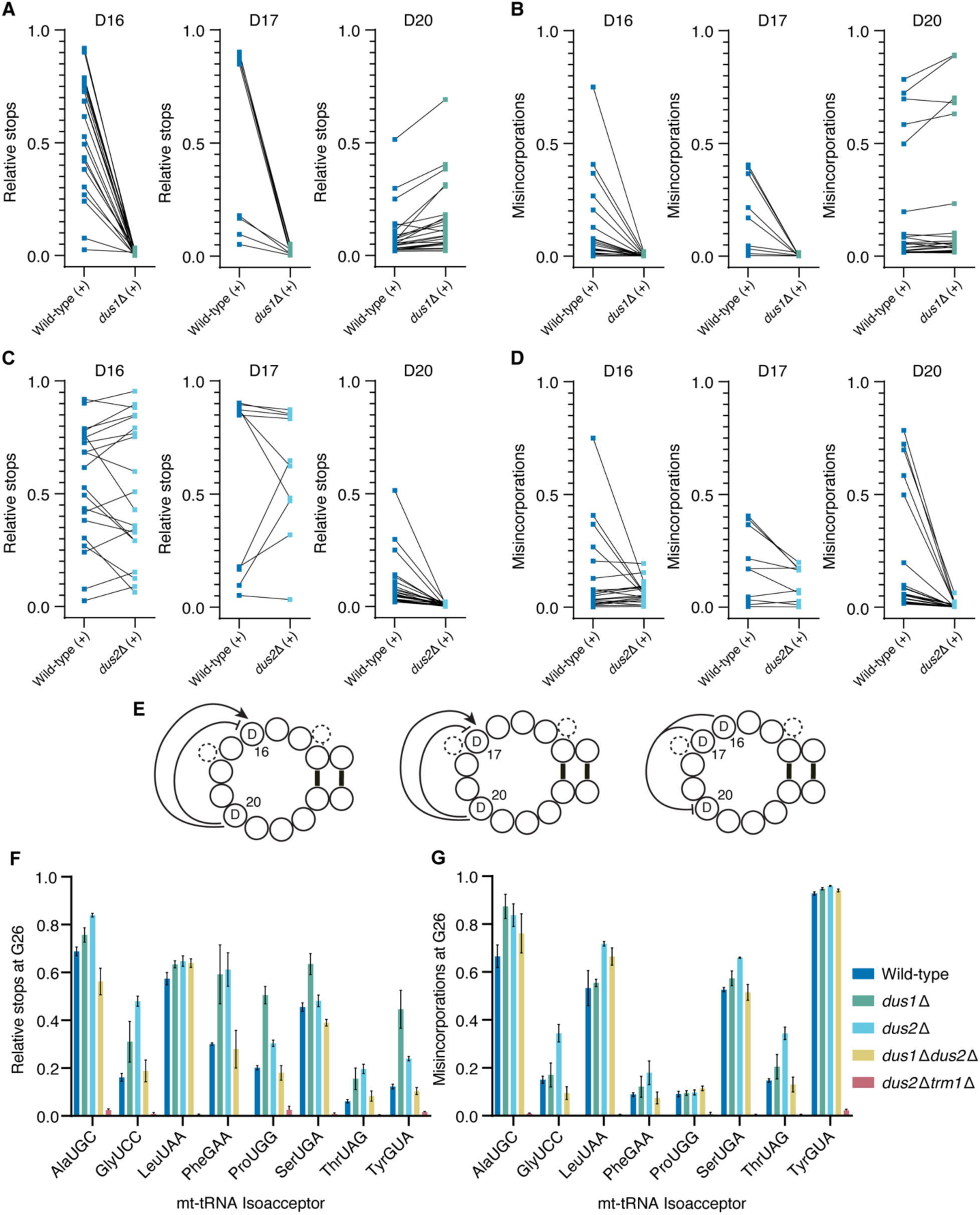
D sites in mt-tRNAs impact the modification levels of other nearby D sites and m^2, 2^G_26_. (**A**, **B**) Relative stops and misincorporations at D-loop positions 16, 17, and 20 in wild-type and *dus1*Δ mt-tRNAs. (**C**, **D**) Relative stops and misincorporations at D-loop positions 16, 17, and 20 in wild-type and *dus2*Δ mt-tRNAs. (**E**) Interactions between dihydrouridines at positions 16, 17, and 20 in mt-tRNAs, based on tD-seq data herein. Arrows represent a positive interaction, while lines ending with a short perpendicular line represent a negative interaction. (**F**, **G**) Levels of m^2, 2^G_26_ in mt-tRNAs upon the loss of Dus1, Dus2, and/or Trm1 activity was determined by relative stops (left) and misincorporations (right) at position G_26_. The mean of three biological replicates is plotted, and the error bars show +/-one standard deviation. All samples included in these plots were treated with NaBH_4_.

### DRS and tD-seq reveal interactions between Dus1, Dus2, and Trm1

The loss of Dus2-catalyzed D_20_ was accompanied by increases or decreases in base miscalls at nearby sites for most mt-tRNAs in our DRS data. For nearly every mt-tRNA with a decrease in miscalls at position 20 in *dus2*Δ, there was an increase (red-colored boxes) in miscall levels at positions 15/16 (Figure 5B). Based on the DRS results, we initially hypothesized that D_20_ represses formation of D_16_/D_17_. We thus used tD-seq to semi-quantitatively measure D levels at Dus1-catalyzed sites in *dus2*Δ. The modification levels at D_16_ and D_17_ generally *decreased* in *dus2*Δ (Figure 7C, D and Supplementary Figure S7B, D), indicating that D_20_ promotes D_16_/D_17_ formation in most mt-tRNAs (Figure 7E, left and middle diagrams), the opposite of our initial DRS-informed hypothesis. However, there were exceptions showing increased modification levels at: D_17a_ for mt-tRNA^Ala(UCG)^, D_17_ for mt-tRNA^Leu(UAA)^ and mt-tRNA^Tyr(GUA)^, and D_14_ for mt-tRNA^Ser(GCU)^ in *dus2*Δ. In these instances, D_20_ *represses* modification by Dus1 (Figure 7E, left and middle diagrams), which suggests that the effect D_20_ has on other D sites depends on the isoacceptor. Since tD-seq shows D_16_/D_17_ levels were not generally increased in *dus2*Δ, the cause of the broadly increased miscalls at positions 15 and 16 in our *dus2*Δ DRS data remains unclear. However, we found a similar pattern in cytosolic tRNAs from the same yeast strain (data not shown). The impact of dihydrouridine on DRS ionic current signal, and therefore the change when it becomes absent, deserves close examination through training with synthetic and biological RNAs. Our use of tD-seq bypassed this concern, however, since it represented a more direct measurement of the dihydrouridine modification than DRS.

In contrast to the variety of outcomes observed upon loss of the *DUS2* gene, loss of *DUS1* led to more consistent changes in Dus2-catalyzed modification levels. The *dus1Δ* mutant showed increased modification levels at D_20_ for several mt-tRNAs in the tD-seq dataset (Figure 7A, B and Supplementary Figure S7A, C). This indicates that Dus1-catalyzed D_14_/D_16_/D_17_/D_17a_ broadly *represses* D_20_ formation in mt-tRNAs (Figure 7E, right diagram).

Several isoacceptors in our DRS data had increased mismatch probabilities at position 26 in the absence of D_20_. For mt-tRNA^Phe(GAA)^, mt-tRNA^Thr(UAG)^, and mt-tRNA^Tyr(GUA)^, we hypothesized this was caused by an increase of the annotated m^2, 2^G_26_ modification, catalyzed by Trm1 (57). We applied tD-seq to determine how the loss of *DUS1* and *DUS2* impacts m^2, 2^G_26_ levels, as this modification disrupts canonical G-C base pairing (58, 59) and thus reverse transcription. In wild-type, most tRNAs containing G_26_ had substantially more relative stops and misincorporations relative to a strain lacking the Trm1 enzyme (*dus2*Δ*trm1*Δ) (Figure 7F, G). Based on this we conclude that the signals observed at G_26_ are due to Trm1-catalyzed m^2, 2^G_26_. Although mt-tRNA^Ser(GCU)^ has a G at position 26, we did not find any evidence of a modification at this site, in agreement with the Modomics database. The *dus1*Δ and *dus2*Δ strains had increased m^2, 2^G_26_ levels, as analyzed by tD-seq, for most m^2, 2^G_26_-containing isoacceptors. This confirmed that the increased DRS-based miscalls at G_26_ for several isoacceptors in *dus2*Δ (Figure 5B) were due to increased m^2, 2^G_26_ levels. While Dus1 does not modify mt-tRNA^Pro(UGG)^, this isoacceptor had elevated levels of m^2, 2^G_26_ in the *dus1*Δ strain (Figure 7F, G). This observation suggests that interactions between Dus1 and Trm1 may be independent from Dus catalytic activity but dependent on Dus binding to mt-tRNAs. In fact, recent findings show that catalytic activity of a Dus1 ortholog, DusB, is not required for its role in an oxidative stress response in *Vibrio cholerae* (60). Surprisingly, the *dus1*Δ*dus2*Δ mutant had m^2, 2^G_26_ levels comparable to wild-type (Figure 7F, G). Collectively these results suggest complex interplay between Dus enzymes and Trm1 that may involve non-catalytic Dus activities. Lastly, mt-tRNA^Lys(UUU)^ showed a change at position 26 in our *dus2*Δ DRS data, however this tRNA contains an adenosine at position 26, and Trm1 is not known or predicted to methylate adenosine bases. Moreover, there was no disruption of RT at this site in our tD-seq data, leaving the source of this signal unclear, necessitating further study to determine if it is a bonafide modification that changes upon loss of Dus2 activity.

## DISCUSSION

As direct RNA sequencing is increasingly applied to study tRNAs from a variety of species and cell-types, it remains important to define reference sequences for each case, while allowing for the *de novo* discovery of new sequences that may be captured. After we updated and expanded the current list of aligned mt-tRNA sequences from *S. cerevisiae*, we used this reference to profile chemical modifications using DRS, identifying the positions that are modified by three enzymes: Pus4, Pus2 and Dus2. We developed tD-seq, a method to map dihydrouridine sites in tRNAs, which we applied to determine all modification sites of Dus1 and Dus2 in mt-tRNAs. In total our study provides a comprehensive map of *S. cerevisiae* mitochondrial tRNA modifications.

Our results revealed novel examples of modification circuits in mt-tRNAs, in which the presence of one modification either stimulates or represses modifications at other sites. In yeast cytosolic tRNAs, the disruption of a single Dus enzyme activity does not lead to changes at other D sites (47). However, here we show that Dus1 represses D_20_ formation and Dus2 can either promote or repress D_14_/D_16_/D_17_/ D_17a_ formation in mt-tRNAs. While we cannot yet explain why Dus1 and Dus2 influence each other’s activities in mt-tRNAs and not in cytosolic tRNAs, there are several contextual differences. Relative to cytosolic tRNAs, mitochondrial tRNAs have lower GC content (62) and fewer modifications, influencing the stability of their tertiary structures. This could make mt-tRNAs more sensitive to changes in individual modifications, resulting in possible compensatory pathways. Cytosolic tRNAs also contain modifications that are not present in mt-tRNAs, which may interfere with the interplay between Dus1 and Dus2 activities that we observed in mt-tRNAs. We also found that the presence of dihydrouridine generally repressed m^2, 2^G_26_ formation by Trm1 in several mt-tRNAs. The contribution of these interactions to mt-tRNA structural stability, and mitochondrial translation, warrants further investigation.

Recent technical developments to Nanopore direct RNA sequencing hold promise for improved tRNA modification detection. This includes improved sequencing accuracy with RNA004 kits and modification-aware basecallers. However, these modification-specific basecalling models are currently limited by high false positive rates (63), the inability to distinguish between chemical isomers such as m^1^A and m^6^A (64), and are currently only available for four RNA modification types. While these models continue to be developed and improved, miscall-based analysis of DRS data remains a suitable approach for detecting tRNA modifications. Our study establishes that DRS can detect dihydrouridine in tRNA using a genetic mutant of a Dus enzyme. This revealed that D causes a miscall-based signal “window” of three neighboring positions. The majority of annotated D sites were indeed detected by DRS, however we also found false negatives in D sites that were detected by tD-seq and reported in the Modomics database, but not detected by DRS. False positives in DRS data were also noted, such as changes in miscalls upon the loss of Dus2 that were not fully recapitulated by tD-seq. Therefore, as we have stressed previously (20), it remains important to pair DRS with orthogonal methods for the most accurate RNA modification detection.

Mitochondrial tRNAs provide an ideal test case for the advancement of direct tRNA-sequencing because they contain many conserved tRNA modifications, but at a lower density compared to cytosolic tRNAs. Therefore, the miscall or ionic current signatures of chemically modified bases are more isolated from neighboring modifications and can be used to advance DRS-based identification of specific modifications. Additionally, each mitochondrial tRNA molecule is transcribed from a single gene copy in the mitochondrial genome, thus bypassing the requirement for analysis of isodecoders that are common among nuclear-encoded tRNA genes. Our work lays a foundation for this type of systematic analysis.

Our approach could be used to model the effects of human disease-associated mutations in tRNA modification enzymes known to impact mitochondrial function. Most tRNA enzyme functions are conserved from yeast to humans (10). Testing human-derived mutations at orthologous sites in yeast proteins, or replacing the yeast proteins with their human counterparts, will expand our understanding of how these mutations affect mitochondrial tRNA function and translation. Our findings also highlight how reduction or loss of a modification can lead to collateral effects on other modifications, and the ensemble of these changes must be considered in context of the resulting molecular and physiological phenotype.

For example, mutations in the human PUS1 gene — whose encoded protein is alone responsible for modifying both cytoplasmic and mitochondrial tRNAs — have been linked to the disease Myopathy Lactic Acidosis Sideroblastic Anemia (MLASA). It is not known whether the MLASA phenotype is due to reduced modification of cytoplasmic tRNAs, mitochondrial tRNAs, or both (65–68). Human PUS1 has two yeast orthologs— Pus1 and Pus2 — that modify cytoplasmic and mitochondrial tRNAs, respectively. Our data showed that loss of Pus2-catalyzed pseudouridylation led to changes in miscalls at other modification sites, consistent with a circuit. Analysis of human cells lacking PUS1 activity will be required to determine whether this also leads to changes in other modifications besides pseudouridine in human mt-tRNAs, as shown here for yeast. If so, this may contribute to the disease phenotype. DRS presents an approach that could be used to address this unknown.

## MATERIALS AND METHODS

### Yeast growth conditions

See Supplementary Table S7 for yeast strains used in this study and their genotypes. Yeast strains were streaked out from glycerol stocks in the -80°C freezer onto YPD plates and grown for 2 days at 30°C. Individual colonies were picked to inoculate 10 mL culture tubes containing 5 mL of YPD media. The liquid cultures were grown overnight on a roller drum wheel at 30°C. The next morning the cultures were diluted 1000-fold into 50 mL of YPD media in 250 mL Erlenmeyer flasks. The cultures were grown overnight on a shaking incubator at 300 rpm and 30°C. The yeast were centrifuged in 50 mL conical tubes at 3000g for 5 min at room temperature. The supernatant was removed, and the pellets were resuspended in 20 mL YPE (Yeast extract, Peptone, 2% Ethanol) media and added to 980 mL YPE in 3 L Erlenmeyer flasks with a baffled base for increased aeration. The cultures were grown on a shaking incubator at 300 rpm and 30°C for ∼36 hours, until an OD_600_ of 1.0-1.4 was reached. The yeast were centrifuged in 1000 ml Nalgene centrifuge bottles at 3000g for 5 min at room temperature. The supernatant was poured off, and the yeast pellets were resuspended in 250 mL of sterile Millipore water and centrifuged again at 3000g for 5 min at room temperature. The supernatant was removed, and the cells were resuspended in 45 mL of 1X PBS (Phosphate Buffered Saline), then transferred to 50 mL conical tubes. The resuspended cells were centrifuged at 3000g and 4°C for 5 min and the supernatant was decanted. The pellets/conical tubes were immediately submerged into liquid nitrogen to flash freeze, then stored at -80°C.

For cells grown in galactose, yeast strains were streaked out from glycerol stocks in the -80°C freezer onto YPD plates and grown for 2 days at 30°C. Individual colonies were picked to inoculate 10 mL culture tubes containing 5 mL of YPD media. The liquid cultures were grown overnight on a roller drum wheel at 30°C. The next morning the cultures were diluted in 5 mL YP 2% galactose media to an OD_600_ of 0.1. The cultures were grown for ∼6 hours at 30°C on a roller drum wheel to an OD_600_ of ∼0.8. The yeast was centrifuged in 15 mL conical tubes at 3000g and 4°C for 2 minutes. The supernatant was removed, and the yeast was resuspended in 1 mL of cold PBS and transferred to microcentrifuge tubes. The tubes were then centrifuged at 9500 RPM for 1 min at 4°C. The supernatant was poured out, and the microcentrifuge tubes were immediately placed into liquid nitrogen to flash freeze yeast in log phase. Total RNA was isolated from the cells as described below, but without mitochondrial enrichment.

### Crude mitochondrial fraction isolation

The crude mitochondrial fraction was isolated from yeast cells using differential centrifugation as previously described (69, 70). The yeast pellets were thawed on ice, resuspended in 45 mL water and centrifuged at 3000g at room temperature for 5 min. The supernatant was decanted, and the wet weight of each cell pellet was recorded. The cell pellets were resuspended in 2 mL of DTT buffer (100 mM Tris-H_2_SO_4_ (pH 9.4), 10 mM dithiothreitol) per gram (wet weight) of cells. The tubes were placed on a shaker at 70 rpm and 30°C for 20 min. The cells were centrifuged at 3000g, room temperature for 5 min. The cell pellets were resuspended in 7 mL Zymolyase buffer (20 mM potassium phosphate (pH 7.4), 1.2 M sorbitol), without Zymolyase, per gram of cells. The cells were centrifuged (3000g, 5 min, 20°C), and the supernatant was decanted. The cells were resuspended in the same volume (7 mL/g cells) of Zymolyase buffer and transferred to a 250 mL Erlenmeyer flask. 1 mg of Zymolyase 100T powder (United States Biological Corporation) per gram of cells was added to the cell suspension. The flasks were placed on a shaker at 70 rpm for 30 min at 30°C. The resulting mixture, which contained spheroplasts from cell wall digestion by Zymolyase, was transferred to 50 mL conical tubes and centrifuged (2200g, 8 min, 4°C). The supernatant was decanted. The pellet was carefully resuspended in 6.5 mL ice-cold homogenization buffer (10 mM Tris-HCl (pH 7.4), 0.6 M sorbitol, 1 mM EDTA, 0.2% (w/v) BSA) per gram of cells. The spheroplasts mixture was centrifuged (2200g, 8 min, 4°C) and the supernatant was decanted. The same volume of homogenization buffer (6.5 mL/g of cells) was used to resuspend the spheroplasts mixture, which was then transferred to a pre-chilled, dounce glass homogenizer on ice. With a tight (“B”) pestle, the spheroplasts mixture was homogenized with 15 strokes. An equal volume of ice-cold homogenization buffer was added to the homogenizer, and then the homogenate was transferred to 50 mL conical tubes on ice. The homogenate was centrifuged (1500g, 5 min, 4°C) to pellet undigested cells, nuclei, and other large debris. The supernatant was poured into a new 50 mL conical tube and centrifuged (3000g, 5 min, 4°C) to further clarify the desired material. The resulting supernatant was poured into a new 50 mL conical tube and centrifuged (12, 000g, 15 min, 4°C) to pellet the mitochondria. The supernatant was decanted, and the pellet was resuspended in 6.5 mL ice-cold homogenization buffer per gram of cells, using trimmed pipette tips to avoid breaking organelles during resuspension. The two previous centrifugation steps (3000g, 5 min, 4°C and 12, 000g, 15 min, 4°C) were repeated once to remove any remaining large debris, and to re-pellet the mitochondria. The supernatant was decanted and the crude mitochondrial fraction (also containing organelles including the endoplasmic reticulum, Golgi, and vacuoles (69)) was resuspended in 3 mL ice-cold SEM buffer (10 mM MOPS-KOH (pH 7.2), 250 mM sucrose, 1 mM EDTA). The crude mitochondrial fraction samples were stored overnight at 4°C in SEM.

### tRNA isolation

The crude mitochondrial fractions were transferred to microcentrifuge tubes and centrifuged (12, 000g, 15 min, 4°C) to pellet the mitochondria. The supernatant was removed, and the mitochondria were resuspended with 400 μL TES buffer (10 mM Tris (pH 7.5), 10 mM EDTA, 0.5% SDS). 400 μL of acidic phenol was added, and the tubes were incubated in a water bath at 65°C for 60 min, with 10 sec of vortexing every 15 min to lyse the mitochondria. The lysates were placed on ice for 5 min, then centrifuged (13, 000 rpm, 10 min, 4°C). The aqueous layer (upper layer) was transferred into a new microcentrifuge tube. 400 μL of chloroform was added and the tube was vortexed for 10 sec. The solution was centrifuged again (13, 000 rpm, 10 min, 4°C), and the chloroform extraction was repeated. The aqueous layer was transferred into a clean microcentrifuge tube, then 1/10 volume of 3M sodium acetate (pH 5.5) and 2.5 volumes of cold 100% EtOH were added. The tubes were mixed by inversion and placed at -80°C for ≥ 1 hr. The tubes were centrifuged (13, 000 rpm, 10 min, 4°C) and the supernatant was removed. 500 μL of cold 70% EtOH was added, and the tubes were once again centrifuged (13, 000 rpm, 10 min, 4°C). The supernatant was removed, and the tubes were left open for 5 min to evaporate remaining EtOH. The pellet was resuspended in 20 μL of water and stored at -80°C.

Using the SequaGel 19:1 Denaturing Gel System (National Diagnostics), an 8% TBE-Urea gel was cast. The gel was pre-run at 45 mA for 30 min. Between 21 and 100 μg of the mitochondrially-enriched RNA was combined with an equal volume of 2X RNA Loading Dye (NEB) and incubated at 70°C for 8 min to denature RNA secondary structure. The samples were loaded onto the gel, which ran at 60 mA for 1 hr in 1X TBE (pH 8.3). The gel was removed from the apparatus, stained with SYBR Gold (Invitrogen) for 10 min, and imaged with an Amersham Typhoon (Cytiva). The tRNA was detected by UV-shadowing and gel segments containing RNA ∼70–100 nt in length were excised. The gel slices were placed into microcentrifuge tubes containing 450 μL of 0.3 M NaCl and incubated overnight at 4°C on a tube inverter. The next day, the liquid was transferred to a new microcentrifuge tube and 1.05 mL of 100% EtOH was added. The samples were incubated at -80°C for ≥ 1 h and centrifuged (13, 000 rpm, 30 min, 4°C). The supernatant was removed, and the pellet was air dried for 5 min to evaporate remaining EtOH. The pellet was resuspended in 10 μL of water, the RNA concentration was measured using the Qubit RNA BR assay kit (Thermo Fisher Scientific), and the tRNA was stored at -80°C.

tRNA used in APM-PAGE experiments was isolated using Nucleobond AX-100 columns (MachereyNagel) as previously described (71).

### Nanopore mt-tRNA sequencing

The RNA and DNA oligonucleotides for the splint adapters were ordered from IDT. Separate 10 μM (7.5 μM for ACCA) stock solutions of the four double-stranded splint adapters were assembled in 1X TNE (10 mM Tris, 50 mM NaCl, 1 mM EDTA) by combining equimolar concentrations of the common adapter that ligates to the 3′ end of the tRNA, and one of four adapter strands complementary to the 3′ tRNA overhang of ACCA, GCCA, UCCA, or CCA (see Supplementary Table S2 for sequences). The splints were annealed by heating at 75°C for 1 min and then cooling to room temperature by leaving the tubes on the bench for 10 min before placing them on ice.

The library preparation was performed as previously described (20), with a few modifications. Briefly, 250 ng of mitochondrial-enriched tRNA was ligated to four different double-stranded splint adapters specific to ACCA (12 pmol), GCCA (8 pmol), UCCA (8 pmol), and CCA (4 pmol) tRNA 3′ overhangs (see Supplementary Table S2 for oligonucleotide sequences). The splint adapter cognate for the CCCA 3′ overhang was excluded as no mitochondrial tRNAs in our reference contain this overhang (Supplementary Table S1). In the first ligation, the splint adapters were ligated to tRNA by T4 RNA Ligase 2 (NEB, stock concentration 10, 000 U/ml, a total of 10 units per reaction). The second ligation of the ONT RMX motor adapter to the first ligation product was performed with T4 DNA ligase (NEB, 2, 000, 000 units/ml). The first and second ligation reaction were each purified with magnetic beads (RNA Clean XP, Beckman Coulter), using 1.8X and 1.5X volume of beads added to the reactions, respectively. The elution of the library, flow cell priming and loading followed the SQK-RNA002 protocol. All sequencing was done on the MinION platform with live basecalling turned off to circumvent MinKnow from discarding reads approximating the length of tRNA reads.

For samples grown in galactose, the library preparation was performed exactly as previously described (20).

### IVT mt-tRNA construction and sequencing

The DNA oligonucleotides for the IVT constructs were purchased from IDT. Their sequences are shown in Supplementary Table S2. Yeast IVT mt-tRNAs were generated using the HiScribe T7 Quick High Yield RNA Synthesis Kit (NEB, E2050), as previously done for yeast cytosolic tRNAs (20). 500 ng of RNA from the IVT reaction was sequenced following the SQK-RNA002 protocol for mRNA sequencing.

### Bioinformatic methods for DRS

Analysis of Nanopore DRS data was done essentially as described (20). Basecalling of ionic current files was performed with Guppy v3.0.3, and the resulting FASTQ files were aligned to a reference — containing all 24 mitochondrial and 42 cytosolic tRNA isoacceptors — using BWA-MEM (parameters ‘-W 13 -k 6 -x ont2d’) (30). Alignments were filtered using SAMtools (72) to retain those with a mapping quality score greater than one (Q1). marginAlign and the subprogram marginCaller (33) were used to generate alignment and error models, and calculate posterior probabilities, respectively. The IVT alignment model, which was derived from a pooled dataset containing mitochondrial IVT tRNA reads (this study) and cytosolic IVT tRNA reads (20), was used as the ‘error model’ in marginCaller in all experiments. Heatmaps containing the posterior probabilities were generated using matplotlib (73).

Mismatch probability (P(A’)), or the probability of any alternative nucleotide call, is complementary to the reference match probability (P(A)).

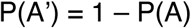

Random subsets of 105 Q1 aligned reads per mt-tRNA isoacceptor were ran through marginCaller and used to calculate mismatch probabilities for all experiments. Miscalls were assigned to positions with a mismatch probability of ≥ 0.3 for biologically derived mt-tRNA samples. To determine novel Pus2-dependent Ψ sites, positions U_27_/U_28_ with a mismatch probability ≥ 0.3 in wild-type and < 0.3 in *pus2*Δ were considered. Changes in mismatch probability (P(A’)_Δ_) were considered significant if the difference was larger than an empirically determined minimum difference threshold (P(A’)_Δ, min_).

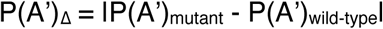

The minimum difference threshold was calculated by taking the mean P(A’)_Δ_ value from the 7 known Ψ_27_/Ψ_28_ sites (8) and subtracting 1 standard deviation. The value of this threshold for Pus2 sites was 0.140.

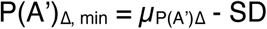

For Dus2 sites, a three-position window centered on U_20_ was examined, but only the position with the largest P(A’)_Δ_ for a given isoacceptor was used in proceeding steps. The minimum difference threshold was calculated as described above from the 16 annotated D_20_ sites and was determined to be 0.223.

### tD-seq (tRNA Dihydrouridine sequencing)

Total RNA isolated from crude mitochondrial fractions was treated with NaBH_4_ as previously described (49). For each sample, 2 μL of 100 mg/mL NaBH_4_ in 10 mM KOH and 1.6 μL of 500 mM Tris-HCl pH 7.5 was added to 1 μg of total RNA in 16.4 μl of water. No-treatment control samples were prepared similarly except for the omission of NaBH_4_ in the 2 μL of 10 mM KOH. Reactions were mixed by pipetting and incubated on ice for 1 hr, then neutralized with 4 μL of 6N acetic acid. RNA was precipitated by adding 1/10 volume of 3M NaAc (pH 5.5), 2.5 volumes of cold 100% ethanol, and 2 μL RNA grade glycogen (Thermo Fisher Scientific) to the neutralized reactions, then placed at -80°C for ≥ 1 hr. The samples were centrifuged (13, 000 rpm, 30 min, 4°C) and the supernatant was removed. The pellets were washed with cold 70% ethanol, and the tubes were once again centrifuged (13, 000 rpm, 10 min, 4°C). The supernatant was removed, and the tubes were left open for 5 min to evaporate remaining ethanol. The RNA pellet was resuspended in 7.8 μL of water.

Ligation of the 3′ reverse transcription adapter was performed as described in the “Sequence-specific Direct RNA protocol” (Oxford Nanopore Technologies) to capture NCCA 3′ tRNA overhangs. Oligo A (RTA A) was ordered without modification, while custom versions of Oligo B (RTA B NCCA) that complement ACCA, UCCA, GCCA, and CCA tRNA ends were used (see Supplementary Table S2 for oligonucleotide sequences). RTA A and RTA B NCCA oligos (1:1) were separately annealed in buffer (10 mM Tris-HCl pH 7.5, 50 mM NaCl) by heating at 95°C for 2 min in a thermocycler, then slowly cooled (0.1°C/sec) to 4°C and placed on ice. The following was added to the 7.8 μL of RNA: 3 μL NEBNext Quick Ligation Reaction Buffer, 0.9 μL ACCA RTA (10 μM), 0.6 μL UCCA RTA (10 μM), 0.6 μL GCCA RTA (10 μM), 0.6 μL CCA RTA (5 μM), and 1.5 μL T4 DNA Ligase (NEB, 2M U/mL). Ligation reactions were mixed by pipetting, incubated at room temperature for 10 min, and immediately used in the following step without cleanup. Reverse transcription was started by adding 9 μL water, 2 μL 10 mM dNTP solution, 8 μL 5X First Strand Buffer, 4 μL 0.1 M DTT, and 2 μL SuperScript III (Invitrogen, 18080093) to the ligation reaction. Reactions were incubated in a thermocycler at 50°C for 50 min, then 70°C for 10 min. RNA was degraded by adding 4 μL 1M NaOH and boiling at 95°C for 5 min, then 4 μL 1M HCl was added to neutralize the solution. The resulting cDNA was ethanol precipitated as described above and resuspended in 8 μL water.

Precipitated cDNA was mixed with an equal volume of 2X RNA loading dye, then heated at 65°C for 5 min and loaded onto an 8% Urea-PAGE gel (8×8 cm) which had been pre-run for 20 min at 200V. The gel was run at 200V in 1X TBE for ∼45 min until the bromophenol blue dye was at the bottom of the gel, then stained with SYBR Gold (1X final concentration in 1X TBE) for 5 min with gentle rocking. Bands corresponding to full length and truncated RT products from tRNAs (∼70-140 nt) were excised and placed in tubes containing 400 μL DNA elution buffer (300 mM NaCl, 10 mM Tris, pH 8.0), then incubated overnight at 4°C on a tube inverter. The eluted cDNA was ethanol precipitated and resuspended in 5 μL of water. Next, 0.8 μL of 3′ adapter (80 μM) and 1 μL DMSO was added to the cDNA, incubated at 75°C for 2 min, and placed on ice for 2 min. The ligation reaction was started by adding 2 μL RNA ligase buffer, 0.2 μL 0.1M ATP, 6.5 μL 50% PEG-8000, 3.6 μL water, and 0.5 μL T4 RNA Ligase 1 (NEB, 10, 000 units/ml), then incubated overnight at 22°C. The reaction was cleaned up with 1.8X volume of magnetic beads (RNA Clean XP, Beckman Coulter) following manufacturer instructions and eluted in 10 μL water. Q5 High-Fidelity DNA Polymerase (NEB, M0491S) was used for library PCR in a final reaction volume of 50 μL, with the following thermocycler program: (1) 98°C 30 sec, (2) 98°C 10 sec, (3) 64°C 30 sec, (4) 72°C 20 sec, (5) 72°C 2 min. Steps 2–4 were repeated for 6 cycles. Excess primers were removed by magnetic bead cleanup with 1.8X volume Mag-Bind TotalPure NGS beads (Omega Bio-tek) according to manufacturer instructions and eluted in 30 μL water. PCR products with unique i5 and i7 indices were pooled and sequenced on the Illumina MiSeq v3 platform using a custom index 1 primer.

Demultiplexed reads were processed as previously described (49), including: trimming of adapter sequences, PCR duplicate collapsing, UMI removal, read mapping, and 5′ read end position gathering. The same tRNA reference library was used for mapping reads from both the DRS and tD-seq datasets, but for tD-seq the RNA bases of the 5′and 3′ DRS splint adapters were not included. Sequencing and alignment statistics for tD-seq Illumina sequencing data are shown in Supplementary Table S9.

Thresholds for D-site calling were empirically determined by examining the ‘relative stops’ and misincorporations at D and non-D uridine (including other modified uridine) sites annotated in the Modomics database. For a position p, the ‘relative stops’ was calculated from the number of mapped reads ending at p + 1 (i.e. not covering p) relative to the number of mapped reads covering p. The difference in relative stops and misincorporations was calculated between matched wild-type biological replicates that were either treated or not treated with sodium borohydride. These differences were averaged across three biological replicates to determine the “mean change in misincorporations” and “mean change in relative stops” for every uridine/modified uridine in the Modomics database. Each site was classified as a “D” or “non-D U” and then ROC curve analysis was performed to determine the optimal thresholds for calling a D-site based on the “mean change in misincorporations” or “mean change in relative stops” (Supplementary Figure S8A). The distribution of these sites relative to the thresholds is plotted in Supplementary Figure S8B. Next, these thresholds were used to identify D-sites in all mt-tRNAs. If a site was over at least one threshold in WT and under both thresholds in *dus1*Δ*dus2*Δ, it was categorized as a D. Positions 1–13 of all mt-tRNA isoacceptors were excluded due to low sequencing coverage at the 5′end. A site was determined to be catalyzed by Dus1 or Dus2 if it was above one threshold in wild-type and below both thresholds in the single deletions (*dus1*Δ or *dus2*Δ).

### APM-PAGE and Northern Blotting

A denaturing gel containing 8% polyacrylamide, 7.5 M urea, and 0.05% ([*N*-acryloylamino]phenyl)mercuric chloride (APM) was cast and pre-ran at 200V for 15 min in 0.5X TBE. Column-isolated, mitochondria-enriched tRNA samples (50 ng) were mixed with an equal volume of 2X RNA Loading Dye (NEB) and heated at 80°C for 5 min. Samples were loaded onto the gel, which ran at 200V until the lower dye front ran off. The RNA was transferred onto a neutral nylon membrane (GVS) in a Trans-Blot Turbo Transfer System (Bio-Rad) for 35 min at 400 mA with 0.5X TBE. RNA was crosslinked to the membrane at 0.12 J/cm^2^ in a UV Stratalinker 1800 (Stratagene). Blots were pre-hybridized in 12.5 mL hybridization buffer (5X SSC, 0.02 M Na_2_HPO_4_ (pH 7.2), 7% SDS, 2X Denhardt) for 4 hr at 50°C with 500 μg sheared salmon sperm DNA (ThermoFisher Scientific). The hybridization buffer and sheared salmon sperm DNA was replaced, and 50 pmol of IR-800 labeled DNA probe (IDT) specific to mt-tRNA^Lys(UUU)^ (sequence from (12)) was added. Blots were hybridized in the dark overnight at 50°C, then washed twice (for 10 min and 30 min) in wash buffer (3X SSC, 0.02 M NaH_2_PO_4_ (pH7.5), 5% SDS, 10X Denhardt), and once in stringent wash buffer (1X SSC, 10% SDS) for 8 min. IR-800 signals were collected using an Amersham Typhoon (Cytiva) and densitometry was performed in Fiji-ImageJ2 (The Fiji Project).

## ACKNOWLEDGEMENTS

We thank Eric Westhof (Université de Strasbourg) for his generous input on optimizing the mt-tRNA structural alignment. We thank Sebastian Leidel (University of Bern) for his sharing and guidance on the tRNA Northern Blot protocol. We thank Ethan Shaw and Hannah Wilson (University of Oregon) for their initial pilot experiment to test enrichment of mt-tRNAs, and Ethan for assistance with the galactose samples. We thank Rosalind Carrier and Margarita Rojas (University of Oregon) for their assistance with Northern Blots. We thank Doug Turnbull and Jeff Bishop (Genomics and Cell Characterization Core Facility, University of Oregon) for their guidance on Illumina sequencing. We thank Charlie Boone (University of Toronto) for the gift of the *dus1*Δ and *dus2*Δ*trm1*Δ yeast strains. We thank Alice Barkan (University of Oregon) and members of the Garcia Lab for comments on the manuscript.

## AUTHOR CONTRIBUTIONS

[J.L.R.] and [D.M.G]: Conceptualization, Methodology, Visualization, Writing — original draft, Writing — review & editing.

[J.L.R]: Data curation, Formal analysis, Investigation. [D.M.G]: Funding acquisition, Project administration.

## DATA AVAILIBILITY

The sequencing data described in this manuscript are deposited at the European Nucleotide Archive. The study accession number is PRJEB89858.

