## Supplementary Data for "Toward a comprehensive modification landscape of yeast mitochondrial tRNAs using Nanopore direct RNA sequencing and Dihydrouridine sequencing"

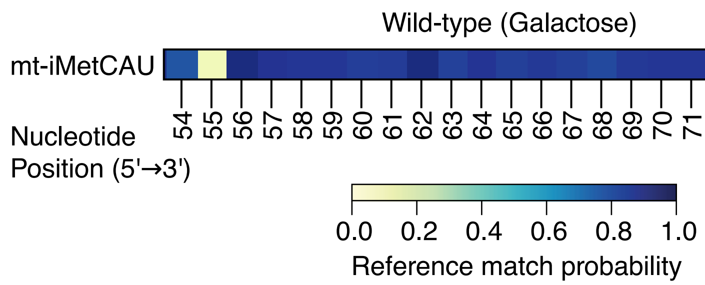


**Supplementary Figure 1.** Heatmap displaying the reference match probabilities of mt-tRNA$^{iMet(CAU)}$ from Wild-type cells grown in YP 2% galactose media. Position 72, which has previously been reported as a pseudouridine (1), is marked with an (*).

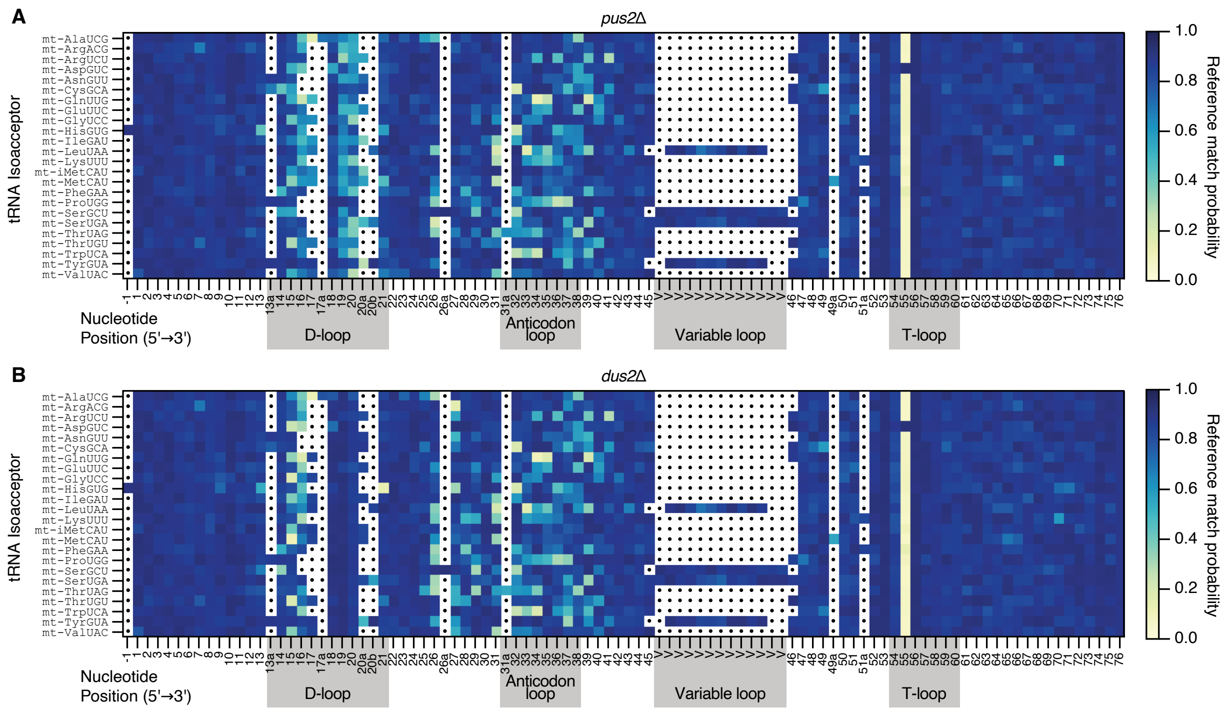


**Supplementary Figure 2.** Heatmaps displaying the reference match probabilities of (**A**) mt-tRNAs from *pus2Δ* cells, and (**B**) mt-tRNAs from *dus2Δ* cells. Otherwise as described in Figure 3.

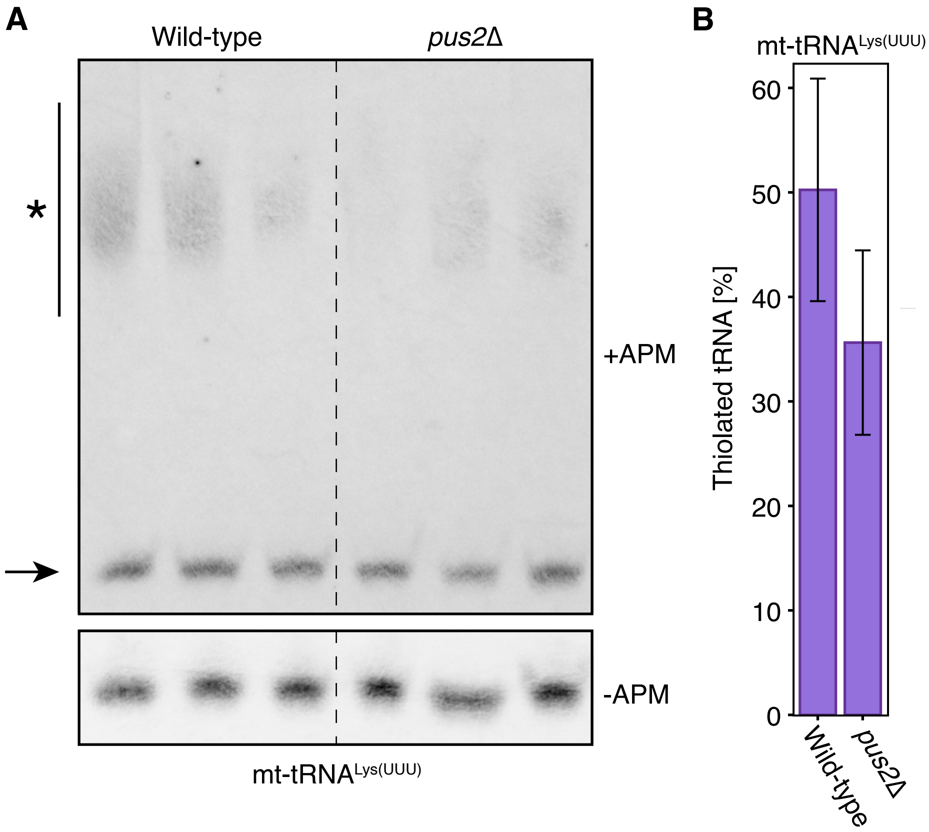


**Supplementary Figure 3.** Analysis of 2-thiouridine levels by APM-PAGE and Northern Blot. (**A**) Mitochondria-enriched tRNAs from wild-type and *pus2*Δ cells were separated on gels either with ([*N*-acryloylamino]phenyl)mercuric chloride (+APM, upper blot) or without APM (-APM, lower blot), transferred to nylon membranes, and probed for mt-tRNA$^{Lys(UUU)}$. The migration of 2-thiolated tRNAs (indicated by an *) was slowed in the +APM gel relative to non-thiolated tRNAs (indicated by an arrow). (**B**) Signals of 2-thiolated and non-thiolated bands were quantified using densitometry, and the percentage of 2-thiolated mt-tRNA$^{Lys(UUU)}$ was calculated. The mean percentage of 2-thiolated mt-tRNA$^{Lys(UUU)}$ was not significantly different between wild-type and *pus2*Δ ($p = 0.2118$, Welch's two-sided T-test).

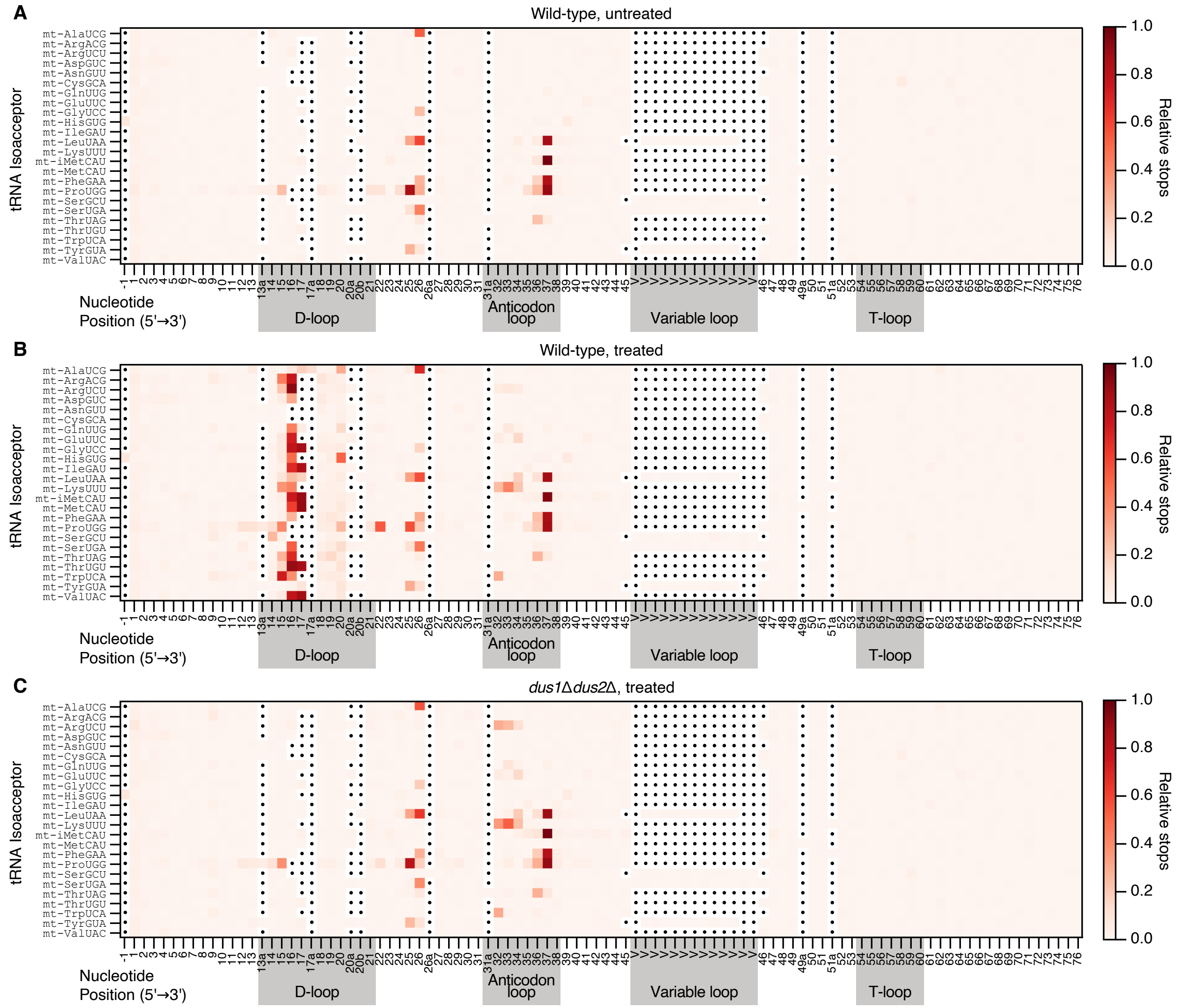


**Supplementary Figure 4.** Heatmaps displaying the average relative stops between three biological replicates from tD-seq. (**A**) Wild-type tRNAs not treated with $NaBH_4$. (**B**) Wild-type tRNAs treated with $NaBH_4$. (**C**) *dus1*Δ*dus2*Δ tRNAs treated with $NaBH_4$.

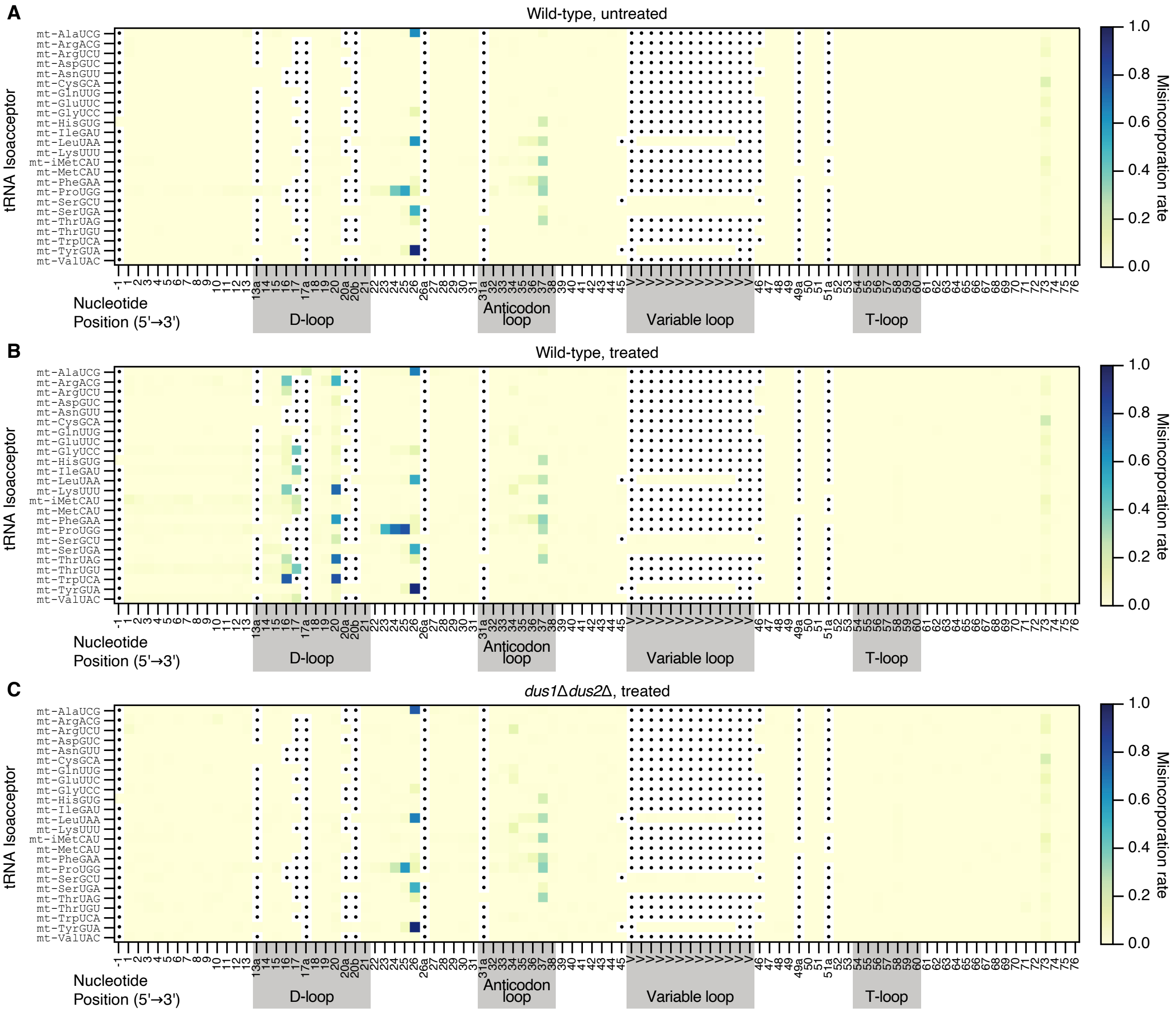


**Supplementary Figure 5.** Heatmaps displaying the average misincorporation rate between three biological replicates from tD-seq. (**A**) Wild-type tRNAs not treated with $NaBH_4$. (**B**) Wild-type tRNAs treated with $NaBH_4$. (**C**) *dus1*Δ*dus2*Δ tRNAs treated with $NaBH_4$.

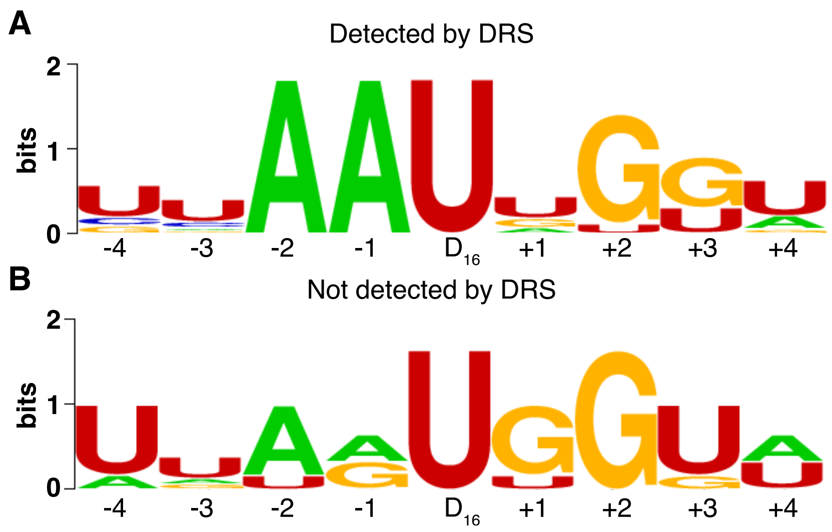


**Supplementary Figure 6.** Sequence logos showing the context of $D_{16}$ sites for (**A**) mt-tRNAs where DRS detected $D_{16}$ and (**B**) mt-tRNAs where DRS did not detect $D_{16}$.

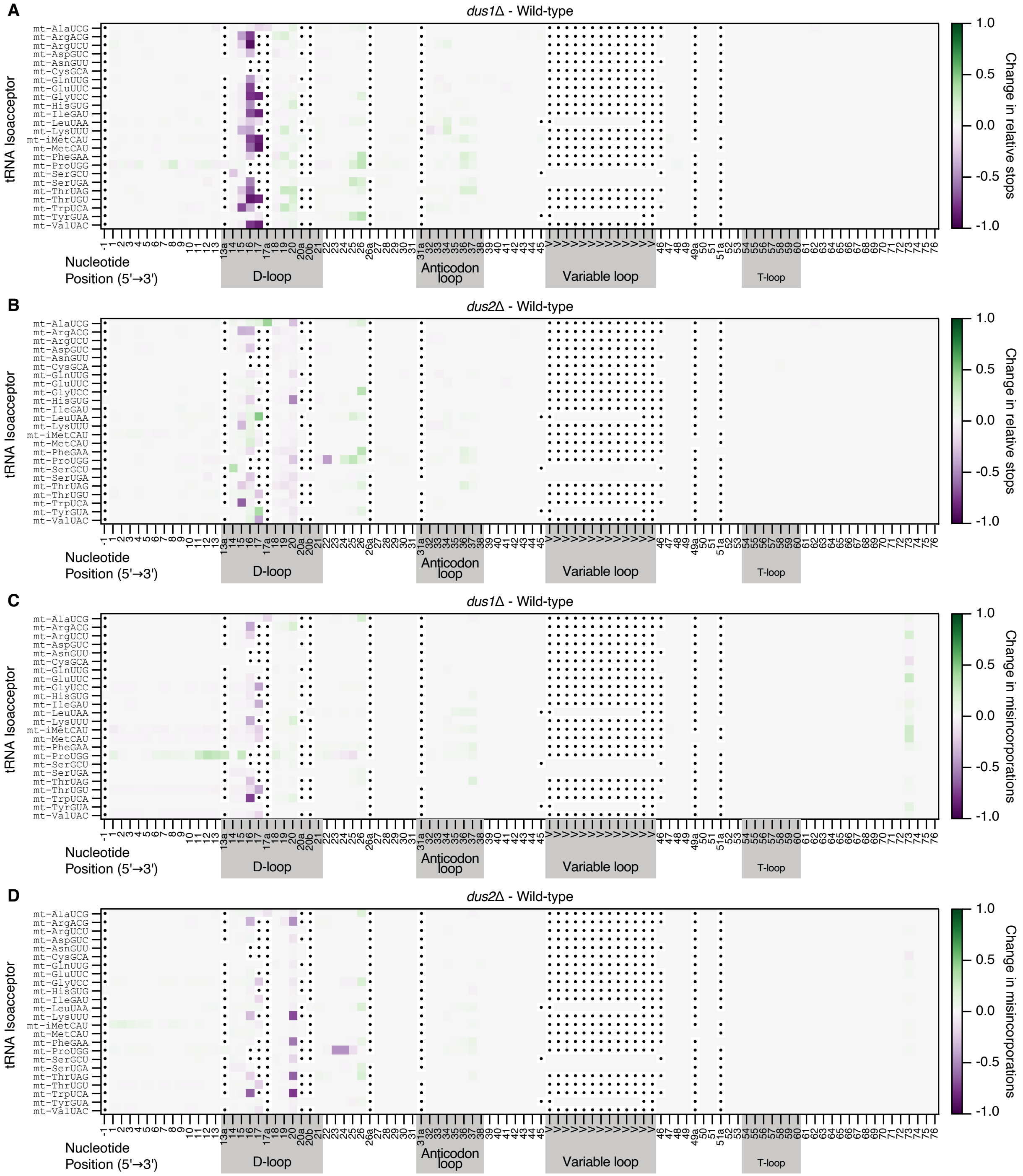


**Supplementary Figure 7.** Heatmaps displaying the change in relative stops (**A**, **B**) and misincorporations (**C**, **D**) between Wild-type and *dus1*Δ (**A**, **C**) or Wild-type and *dus2*Δ (**B**, **D**). All samples displayed were treated with $NaBH_4$.

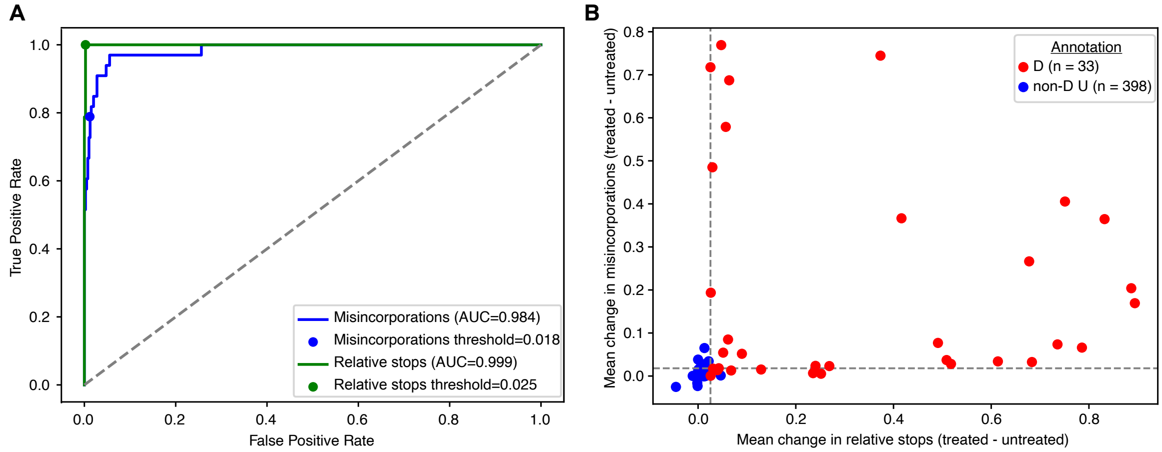


**Supplementary Figure 8.** Threshold determination for D-site calling with tD-seq. Uridines were classified as D or non-D Uridines based on mt-tRNA sequences in Modomics. Sequences not represented in Modomics were not included in this analysis. Additionally, position $U_{34}$ was not included if annotated to be modified. (**A**) Receiver operating curves for D-site classification, based on the mean change in misincorporations (blue line) or relative stops (green line) between matched wild-type samples either treated or not treated with $NaBH_4$. (**B**) Scatter plot showing the distribution of D vs non-D uridine sites relative to the thresholds (grey dotted lines).

**Supplementary Table S1.** Reference sequences for 24 yeast mitochondrial and 42 yeast cytosolic tRNA isoacceptors from strain BY4741. Note that the RNA bases of the 5´ and 3´ splint adapters are included. See Supplementary_Tables_S1_S2.xlsx

**Supplementary Table S2.** The RNA and DNA oligonucleotides for the splint adapters and IVT constructs. See Supplementary_Tables_S1_S2.xlsx

**Supplementary Table S3.** Sequencing and alignment statistics for direct RNA sequencing. Wild-type samples with an * were grown in YP 2% galactose media (see Materials and Methods).

| | **Per flow cell** | | | |
|---|---|---|---|---|
| **Sample** | **Total reads** | **Aligned tRNA reads** | **Aligned mitochondrial tRNA reads** | **Mitochondrial tRNA reads as a percent of aligned tRNA reads** |
| Wild-type (YDG1) N = 2 | 47,354 – 52,950 | 20,587 – 25,442 | 16,479 – 22,630 | 80.0 – 88.9% |
| *pus4*Δ (YDG127) N = 2 | 34,843 – 37,750 | 15,337 – 19,040 | 11,940 – 16,372 | 77.9 – 86.0% |
| *pus2*Δ (YDG893) N = 2 | 66,220 – 97,163 | 36,597 – 52,075 | 32,466 – 40,736 | 88.7 – 78.2% |
| *dus2*Δ (YDG974) N = 2 | 84,242 – 112,800 | 48,210 – 63,709 | 46,847 – 62,516 | 97.2 – 98.1% |
| IVT N = 1 | 541,217 | N/A | 419,116 | N/A |
| Wild-type* (YDG1) N = 3 | 54,091 – 129,457 | 24,553 – 57,659 | 583 – 1,457 | 1.8 – 2.5% |

**Supplementary Table S4.** Agreement of DRS with annotated modifications on Modomics (mismatch probability ≥ 0.3), from wild-type cells (grown in Ethanol). Note that the list of modifications is derived only from the 16 tRNAs represented on Modomics currently (in parentheses next to modification name is the character used for that in Modomics). Mismatch probabilities were calculated using subsampled reads (n=105) from wild-type cells. This number was chosen as it corresponded to the highest number of reads obtained for the lowest expressed tRNA (mt-tRNA$^{Cys(GCA)}$) in our *pus4*Δ dataset, which had the fewest overall reads of all experiments. Percent agreement is calculated based on the number of tRNAs with a specific modification detected by DRS at/near a specific position divided by the total number of mt-tRNAs for which this modification is documented in Modomics.

| Position | Modification | mt-tRNAs on Modomics with modification | mt-tRNAs with modification detected by DRS | Percent agreement |
|---|---|---|---|---|
| 14 | Dihydrouridine (D) | 1 | 1 | 100 |
| 16 | Dihydrouridine (D) | 14 | 8 | 57 |
| 17 | Dihydrouridine (D) | 2 | 2 | 100 |
| 20 | Dihydrouridine (D) | 16 | 13 | 88 |
| 26 | 2N-methylguanosine (L) | 4 | 4 | 100 |
| 26 | N2 -dimethylguanosine (R) | 3 | 2 | 67 |
| 27 | Pseudouridine (P) | 6 | 6 | 100 |
| 28 | Pseudouridine (P) | 1 | 1 | 100 |
| 31 | Pseudouridine (P) | 7 | 7 | 100 |
| 32 | Pseudouridine (P) | 6 | 5 | 83 |
| 34 | Carboxymethylaminomethyluridine (!) | 2 | 2 | 100 |
| 34 | Carboxymethylaminomethyl-2-thiouridine ($) | 1 | 1 | 100 |
| 37 | N6-threonylcarbamoyladenosine (6) | 4 | 3 | 75 |
| 37 | Unknown modified adenosine (H) | 1 | 1 | 100 |
| 37 | 1N-methylguanosine (K) | 7 | 5 | 71 |
| 37 | N6-isopentenyl-adenosine (Ч) | 2 | 1 | 50 |
| 38 | Pseudouridine (P) | 2 | 1 | 50 |
| 39 | Pseudouridine (P) | 6 | 5 | 83 |
| 54 | 5-methyluridine (T) | 15 | 0 | 0 |
| 55 | Pseudouridine (P) | 16 | 16 | 100 |
| 72 | Pseudouridine (P) | 1 | 0 | 0 |

**Supplementary Table S5.** Mismatch probabilities from IVT, wild-type, *pus4*Δ, *pus2*Δ, and *dus2*Δ DRS experiments. Positions with a mismatch probability of ≥ 0.3 are highlighted in yellow. See Supplementary_Table_S5.xlsx

**Supplementary Table S6.** Summary of Pus4, Pus2, and Dus2 dependent modifications in mt-tRNAs determined by DRS and confirmed via genetic mutant. Annotation status based on documentation in the Modomics database (2).

| mt-tRNA isoacceptor | Position | Modification | Catalyzing enzyme | Annotation status |
|---|---|---|---|---|
| ArgACG | 27 | P | Pus2 | known |
| ArgUCU | 27 | P | Pus2 | known |
| AsnGUU | 27 | P | Pus2 | novel |
| HisGUG | 27 | P | Pus2 | known |
| SerUGA | 27 | P | Pus2 | known |
| ThrUGU | 27 | P | Pus2 | novel |
| TyrGUA | 27 | P | Pus2 | known |
| ValUAC | 27 | P | Pus2 | novel |
| LysUUU | 28 | P | Pus2 | known |
| ThrUGU | 28 | P | Pus2 | novel |
| AlaUCG | 55 | P | Pus4 | novel |
| ArgACG | 55 | P | Pus4 | known |
| ArgUCU | 55 | P | Pus4 | known |
| AsnGUU | 55 | P | Pus4 | novel |
| CysGCA | 55 | P | Pus4 | novel |
| GlnUUG | 55 | P | Pus4 | novel |
| GluUUC | 55 | P | Pus4 | novel |
| GlyUCC | 55 | P | Pus4 | known |
| HisGUG | 55 | P | Pus4 | known |
| IleGAU | 55 | P | Pus4 | known |
| LeuUAA | 55 | P | Pus4 | known |
| LysUUU | 55 | P | Pus4 | known |
| iMetCAU | 55 | P | Pus4 | known |
| MetCAU | 55 | P | Pus4 | known |
| PheGAA | 55 | P | Pus4 | known |
| ProUGG | 55 | P | Pus4 | known |
| SerGCU | 55 | P | Pus4 | known |
| SerUGA | 55 | P | Pus4 | known |
| ThrUAG | 55 | P | Pus4 | known |
| ThrUGU | 55 | P | Pus4 | novel |
| TrpUCA | 55 | P | Pus4 | known |
| TyrGUA | 55 | P | Pus4 | known |
| ValUAC | 55 | P | Pus4 | novel |

**Supplementary Table S7.** Yeast strains used in this study.

| Strain | Phenotype | Genotype | Source |
|---|---|---|---|
| YDG1 | Wild-type | MATa, *his3*Δ1, *leu2*Δ0, *met15*Δ0, *ura3*Δ0 | (3) |
| YDG127 | Lacks Ψ55 | MATa, *PUS4*::KanMX, *his3*Δ1, *leu2*Δ0, *met15*Δ0, *ura3*Δ0 | Yeast Knockout Library (4) |
| YDG893 | Lacks Ψ27, Ψ28 | MATa, *PUS2*::KanMX, *his3*Δ1, *leu2*Δ0, *met15*Δ0, *ura3*Δ0 | Yeast Knockout Library (4) |
| YDG974 | Lacks D20 | MATa, *DUS2*::KanMX, *his3*Δ1, *leu2*Δ0, *met15*Δ0, *ura3*Δ0 | Yeast Knockout Library (4) |
| YDG1281 | Lacks D14, D16, D17, D17a, D20 | MATa, *DUS2*::KanMX, *DUS1*::URA3 *his3*Δ1, *leu2*Δ0, *met15*Δ0, *ura3*Δ0 | This study |
| YDG1312 | Lacks D20, $m^{2,2}$G26 | MATα, *TRM1*::natR *DUS2*::KlURA3 *can*Δ::STE2pr-Sp_his5 *lyp1*ΔSTE3pr-LEU2 | Boone Lab, University of Toronto |
| YDG1314 | Lacks D14, D16, D17, D17a | MATα, *dus1*Δ0::natMX4 *can1*Δ0::STE2pr-Sp_HIS5 *lyp1*Δ0 *his3*Δ1 *leu2*Δ0 *ura3*Δ0 *met15*Δ0 LYS2+ | Boone Lab, University of Toronto |

**Supplementary Table S8.** Comparison of modification annotations determined by Modomics, DRS, and tD-seq in wild-type yeast mitochondrial tRNAs. Blue circles indicate/predict presence, magenta circles indicate/predict absence. ND indicates the site is not described in Modomics, and ND$^{!}$ indicates that Modomics incorrectly annotated a site as mono-methylated $m^2G_{26}$; Trm1 is only known to catalyze the formation of di-methylated $m^{2,2}G_{26}$. In the Enzyme (tD-seq) column, (#) represents that the site was under both thresholds in *dus1*Δ, but that there was a change in one threshold in *dus2*Δ as well, (*) represents that the site was over the relative stop threshold in the assigned *dus*Δ strain, and (^) represents that the site was over the misincorporation threshold in the assigned *dus*Δ strain.

| tRNA | Position | Modification | Modomics | DRS | tD-seq | Enzyme (tD-seq) |
|---|---|---|---|---|---|---|
| mt-AlaUGC | 16 | D | 🔵 | 🔵 | 🔵 | Dus1 |
| mt-AlaUGC | 17 | D | ND | 🔵 | 🔵 | Dus1$^{\#}$ |
| mt-AlaUGC | 17a | D | ND | 🔵 | 🔵 | Dus1 |
| mt-AlaUGC | 26 | $m^{2,2}G$ | ND | 🔵 | 🔵 | Trm1 |
| mt-AlaUGC | 20 | D | ND | 🔵 | 🔵 | Dus2 |
| mt-AlaUCG | 62 | unknown | ND | 🔴 | 🔵 | |
| mt-ArgACG | 16 | D | 🔵 | 🔵 | 🔵 | Dus1 |
| mt-ArgACG | 20 | D | 🔵 | 🔵 | 🔵 | Dus2 |
| mt-ArgUCU | 16 | D | 🔵 | 🔴 | 🔵 | Dus1 |
| mt-ArgUCU | 20 | D | 🔵 | 🔵 | 🔵 | Dus2 |
| mt-ArgUCU | 34 | unknown | 🔵 | 🔵 | 🔵 | |
| mt-AsnGUU | 20 | D | ND | 🔵 | 🔵 | Dus2 |
| mt-AspGUC | 16 | D | ND | 🔵 | 🔵 | Dus1 |
| mt-AspGUC | 20 | D | ND | 🔴 | 🔵 | Dus2 |
| mt-CysGCA | 20 | D | ND | 🔴 | 🔵 | Dus2 |
| mt-GlnUUG | 16 | D | ND | 🔵 | 🔵 | Dus1 |
| mt-GlnUUG | 20 | D | ND | 🔵 | 🔵 | Dus2 |
| mt-GlnUUG | 34 | $cmnm^5s^2U$ | ND | 🔵 | 🔵 | |
| mt-GluUUC | 16 | D | ND | 🔵 | 🔵 | Dus1 |
| mt-GluUUC | 20 | D | ND | 🔵 | 🔵 | Dus2 |
| mt-GluUUC | 34 | $cmnm^5s^2U$ | ND | 🔵 | 🔵 | |
| mt-GlyUCC | 16 | D | 🔵 | 🔵 | 🔵 | Dus1 |
| mt-GlyUCC | 17 | D | 🔴 | 🔵 | 🔵 | Dus1 |
| mt-GlyUCC | 20 | D | 🔵 | 🔵 | 🔵 | Dus2 |
| mt-GlyUCC | 26 | $m^{2,2}G$ | 🔵 | 🔴 | 🔵 | Trm1 |
| mt-HisGUG | 16 | D | 🔵 | 🔵 | 🔵 | Dus1 |
| mt-HisGUG | 20 | D | 🔵 | 🔵 | 🔵 | Dus2 |
| mt-IleGAU | 16 | D | 🔵 | 🔵 | 🔵 | Dus1 |
| mt-IleGAU | 17 | D | 🔵 | 🔵 | 🔵 | Dus1 |
| mt-IleGAU | 20 | D | 🔵 | 🔵 | 🔵 | Dus2 |
| mt-iMetCAU | 16 | D | 🔵 | 🔵 | 🔵 | Dus1 |
| mt-iMetCAU | 17 | D | 🔴 | 🔵 | 🔵 | Dus1 |
| mt-iMetCAU | 20 | D | 🔵 | 🔵 | 🔵 | Dus2 |

| tRNA | Position | Modification | Modomics | DRS | tD-seq | Enzyme (tD-seq) |
|---|---|---|---|---|---|---|
| mt-LeuUAA | 16 | D | 🔵 | 🔵 | 🔵 | Dus1 |
| mt-LeuUAA | 17 | D | 🟠 | 🔵 | 🔵 | Dus1* |
| mt-LeuUAA | 20 | D | 🔵 | 🔵 | 🔵 | Dus2 |
| mt-LeuUAA | 26 | $m^{2,2}G$ | 🔵 | 🔵 | 🔵 | Trm1 |
| mt-LeuUAA | 34 | $cmnm^5U$ | 🔵 | 🔵 | 🔵 | |
| mt-MetCAU | 17 | D | 🔵 | 🔵 | 🔵 | Dus1 |
| mt-MetCAU | 20 | D | 🔵 | 🔵 | 🔵 | Dus2 |
| mt-PheGAA | 16 | D | 🔵 | 🟠 | 🔵 | Dus1*/Dus2* |
| mt-PheGAA | 20 | D | 🔵 | 🔵 | 🔵 | Dus2 |
| mt-PheGAA | 26 | $m^{2,2}G$ | 🔵 | 🔵 | 🔵 | Trm1 |
| mt-ProUGG | 20 | D | 🔵 | 🟠 | 🔵 | Dus2 |
| mt-ProUGG | 26 | $m^{2,2}G$ | ND! | 🔵 | 🔵 | Trm1 |
| mt-SerGCU | 14 | D | 🔵 | 🔵 | 🔵 | Dus1 |
| mt-SerGCU | 20 | D | 🔵 | 🔵 | 🔵 | Dus2 |
| mt-SerUGA | 16 | D | 🔵 | 🟠 | 🔵 | Dus1 |
| mt-SerUGA | 20 | D | 🔵 | 🔵 | 🔵 | Dus2 |
| mt-SerUGA | 26 | $m^{2,2}G$ | ND! | 🔵 | 🔵 | Trm1 |
| mt-ThrUAG | 16 | D | 🔵 | 🟠 | 🔵 | Dus1 |
| mt-ThrUAG | 20 | D | 🔵 | 🔵 | 🔵 | Dus2^ |
| mt-ThrUAG | 26 | $m^{2,2}G$ | ND! | 🔵 | 🔵 | Trm1 |
| mt-ThrUGU | 16 | D | ND | 🔵 | 🔵 | Dus1 |
| mt-ThrUGU | 17 | D | ND | 🔵 | 🔵 | Dus1 |
| mt-ThrUGU | 20 | D | ND | 🔵 | 🔵 | Dus2 |
| mt-TrpUCA | 16 | D | 🔵 | 🟠 | 🔵 | Dus1 |
| mt-TrpUCA | 20 | D | 🔵 | 🔵 | 🔵 | Dus2 |
| mt-TyrGUA | 16 | D | 🔵 | 🟠 | 🔵 | Dus1 |
| mt-TyrGUA | 17 | D | 🟠 | 🟠 | 🔵 | Dus1 |
| mt-TyrGUA | 20 | D | 🔵 | 🔵 | 🔵 | Dus2 |
| mt-TyrGUA | 26 | $m^{2,2}G$ | ND! | 🔵 | 🔵 | Trm1 |
| mt-ValUAC | 16 | D | ND | 🔵 | 🔵 | Dus1 |
| mt-ValUAC | 17 | D | ND | 🟠 | 🔵 | Dus1 |
| mt-ValUAC | 20 | D | ND | 🔵 | 🔵 | Dus2 |

**Supplementary Table S9.** Sequencing and alignment statistics for tD-seq Illumina sequencing data.

| Sample | Total reads | Mean Quality Score (PF) | %Q30 | Deduplicated reads | Mapped reads |
|---|---|---|---|---|---|
| WT_rep1_untreated | 183644604 | 34.95 | 0.88 | 1,025,813 | 985744 |
| WT_rep2_untreated | 231148164 | 35.27 | 0.9 | 1,255,003 | 1207081 |
| WT_rep3_untreated | 185003832 | 34.82 | 0.88 | 1,029,360 | 986050 |
| WT_rep1_NaBH4treated | 179897328 | 33.56 | 0.83 | 991,553 | 948718 |
| WT_rep2_NaBH4treated | 275750592 | 33.68 | 0.83 | 1,479,793 | 1417101 |
| WT_rep3_NaBH4treated | 341150784 | 34.27 | 0.85 | 1,851,634 | 1750933 |
| dus2del_rep1_NaBH4treated | 272453532 | 34.17 | 0.85 | 1,479,092 | 1426149 |
| dus2del_rep2_NaBH4treated | 234248352 | 34.76 | 0.87 | 1,271,340 | 1236967 |
| dus2del_rep3_NaBH4treated | 229972548 | 34.06 | 0.85 | 1,262,128 | 1240386 |
| dus1del_dus2del_rep1_NaBH4treated | 196089348 | 34.48 | 0.86 | 1,099,951 | 1055534 |
| dus1del_dus2del_rep2_NaBH4treated | 262902276 | 35.19 | 0.89 | 1,448,600 | 1400365 |
| dus1del_dus2del_rep3_NaBH4treated | 199516980 | 34.51 | 0.87 | 1,128,144 | 1083884 |
| dus2del_trm1del_rep1_NaBH4treated | 178378980 | 34.17 | 0.85 | 1,009,513 | 984099 |
| dus2del_trm1del_rep2_NaBH4treated | 246341472 | 34.36 | 0.86 | 1,366,746 | 1307149 |
| dus2del_trm1del_rep3_NaBH4treated | 334383816 | 34.68 | 0.87 | 1,837,019 | 1732217 |
| dus1del_rep1_NaBH4treated | 241418424 | 34.38 | 0.86 | 1,340,631 | 1307985 |
| dus1del_rep2_NaBH4treated | 229723884 | 34.93 | 0.88 | 1,277,097 | 1252181 |
| dus1del_rep3_NaBH4treated | 208611624 | 34.16 | 0.85 | 1,171,332 | 1148869 |
